# Presynaptic Gq-coupled receptors drive biphasic dopamine transporter trafficking that modulates dopamine clearance and motor function

**DOI:** 10.1101/2021.06.04.447129

**Authors:** Patrick J. Kearney, Elizabeth Kahuno, Tucker L. Conklin, Gilles E. Martin, Gert Lubec, Haley E. Melikian

## Abstract

Extracellular dopamine (DA) levels are constrained by the presynaptic DA transporter (DAT), a major psychostimulant target. Despite its necessity for DA neurotransmission, DAT regulation *in situ* is poorly understood, and it is unknown whether regulated DAT trafficking impacts dopaminergic signaling and/or behaviors. Leveraging chemogenetics and conditional gene silencing, we found that activating presynaptic Gq-coupled receptors, either hM3Dq or mGluR5, drove rapid biphasic DAT membrane trafficking, with region-specific differences in ventral and dorsal striata. DAT insertion required DRD2 autoreceptors and intact retromer, whereas DAT retrieval required PKC activation and Rit2. *Ex vivo* voltammetry revealed that DAT trafficking impacts DA clearance. Importantly, dopaminergic mGluR5 silencing elevated surface DAT, which abolished motor learning and was rescued by inhibiting DAT. We found that presynaptic DAT trafficking is complex, multimodal, and region-specific, and identify cell autonomous mechanisms governing presynaptic DAT tone. Importantly, the findings suggest regulated DAT trafficking impacts both DA clearance and motor function.

## Introduction

Dopamine (DA) is critical for movement, learning, motivation and reward^1, 2^, and DAergic dysfunction is implicated in multiple neuropsychiatric disorders including Parkinson’s Disease (PD), attention-deficit hyperactivity disorder (ADHD), schizophrenia, and addiction^3–6^. Following its release, DA is temporally and spatially constrained by the presynaptic DA transporter (DAT), which recaptures extracellular DA^7^. DAT’s central role in DAergic transmission is illustrated by the consequences of pharmacologically or genetically silencing DAT^8^. For example, DAT is the primary target for addictive and therapeutic psychostimulants, such as cocaine, methylphenidate (Ritalin), and amphetamine (AMPH). These agents markedly enhance extracellular DA through their actions as DAT inhibitors (cocaine, methylphenidate) and substrates (AMPH). Moreover, multiple DAT coding variants have been reported in probands from ADHD, and autism spectrum disorder (ASD) patients^9–14^, as well as in DAT deficiency syndrome, a form of Parkinsonism^15–19^. Given that DAT function profoundly impacts DAergic signaling, it is vital that we understand the molecular mechanisms that acutely regulate DAT availability. Such information may inform potential interventions for DA-related neuropsychiatric disorders.

Extensive research from multiple laboratories supports that DAT surface expression is regulated by membrane trafficking^20–23^. DAT constitutively cycles between the plasma membrane and endosomes. Direct PKC activation accelerates DAT endocytosis, and thereby diminishes DAT surface levels and function^24–26^. PKC-stimulated DAT internalization requires the neuronal GTPase Rit2^27^, whereas DAT endocytic recycling requires the Vps35^+^ retromer complex^28, 29^. In previous studies, PKC was directly activated with phorbol esters or via a Gq-coupled receptor in non-DAergic cell lines. However, it is still unknown whether, or how, DAT traffics in *bona fide* DAergic terminals in response to endogenous, receptor-mediated PKC activation. Further, it is unknown whether regulated DAT trafficking is regionally distinct, or whether DAT trafficking impacts DAergic signaling and DA-dependent behaviors. In this study, we leveraged chemogenetics and *in vivo* conditional gene silencing, complemented by pharmacological, electrochemical and behavioral approaches, to directly probe these questions. Our results demonstrate that regulated DAT trafficking is significantly more complex than had been previous appreciated, and that mechanisms that perturb DAT trafficking also dysregulate DA signaling and DA-dependent motor behaviors.

## Results

### Gq-coupled DREADD activation drives region-specific, biphasic DAT trafficking

Previous DAT trafficking studies activated PKC with phorbol esters, which broadly activate multiple PKC isoforms and do so globally across all cells within heterogenous tissue preparations. Conventional PKCs are most commonly activated by Gq-coupled receptors, and previous studies in non-neuronal cell lines, midbrain, and synaptosomes reported that Gq-coupled receptor activation drives PKC-dependent DAT internalization and/or functional downregulation^30–32^. Given that DAT trafficking may differ in DA neurons vs. cell lines, and that global PKC activation may drive DAT trafficking indirectly within the local striatal circuitry, we first aimed to test whether cell-autonomous presynaptic Gq-coupled signaling impacts DAT surface availability in *bona fide* striatal DAergic terminals. We leveraged the Tet-OFF system to conditionally express the Gq-coupled DREADD, hM3Dq, in *Pitx3^IRES-tTA^;TRE-HA-hM3Dq* mouse DAergic neurons^33^. Pitx3 is a DA-specific transcription factor and previous work from our laboratory and others demonstrated that *Pitx3^IRES-tTA^* selectively drives gene expression in midbrain DA neurons^34, 35^. We first used surface biotinylation to ask whether DAergic hM3Dq activation modulates DAT surface expression in *ex vivo* striatal slices from *Pitx3^IRES-tTA^;TRE-HA-hM3Dq* mice, containing both ventral and dorsal striata. Surprisingly, treatment with the DREADD-specific agonist clozapine-N-oxide (CNO^33^, 500nM, 37°C) biphasically modulated DAT surface expression, which significantly increased by 5 min, and returned to baseline by 30 min (Fig. 1a). Importantly, CNO had no significant effect on DAT surface levels in striatal slices from control *Pitx3^IRES-tTA^/+* littermates (Fig. 1a). We further asked whether Gq-stimulated DAT trafficking was either sex- or region-specific in ventral (VS) and dorsal (DS) striatal subregions. hM3Dq-stimulated DAT trafficking did not significantly differ between males and females (Two-way ANOVA, no effect of sex, p=0.59 (DS), p=0.36 (VS)), therefore data from male and female mice were pooled. Similar to total striatum, DAT was rapidly inserted into the plasma membrane both VS and DS in response to CNO treatment (500nM, 5 min, 37°C, Fig. 1b). In VS, DAT surface expression significantly diminished to baseline levels after 10 min (Fig. 1b). In contrast, in DS, surface DAT remained elevated after 10 min, but returned to baseline by 30 min. DAT surface expression in DS remained significantly higher than VS at both 10 and 30 min (Fig. 1b).

**Figure 1.**
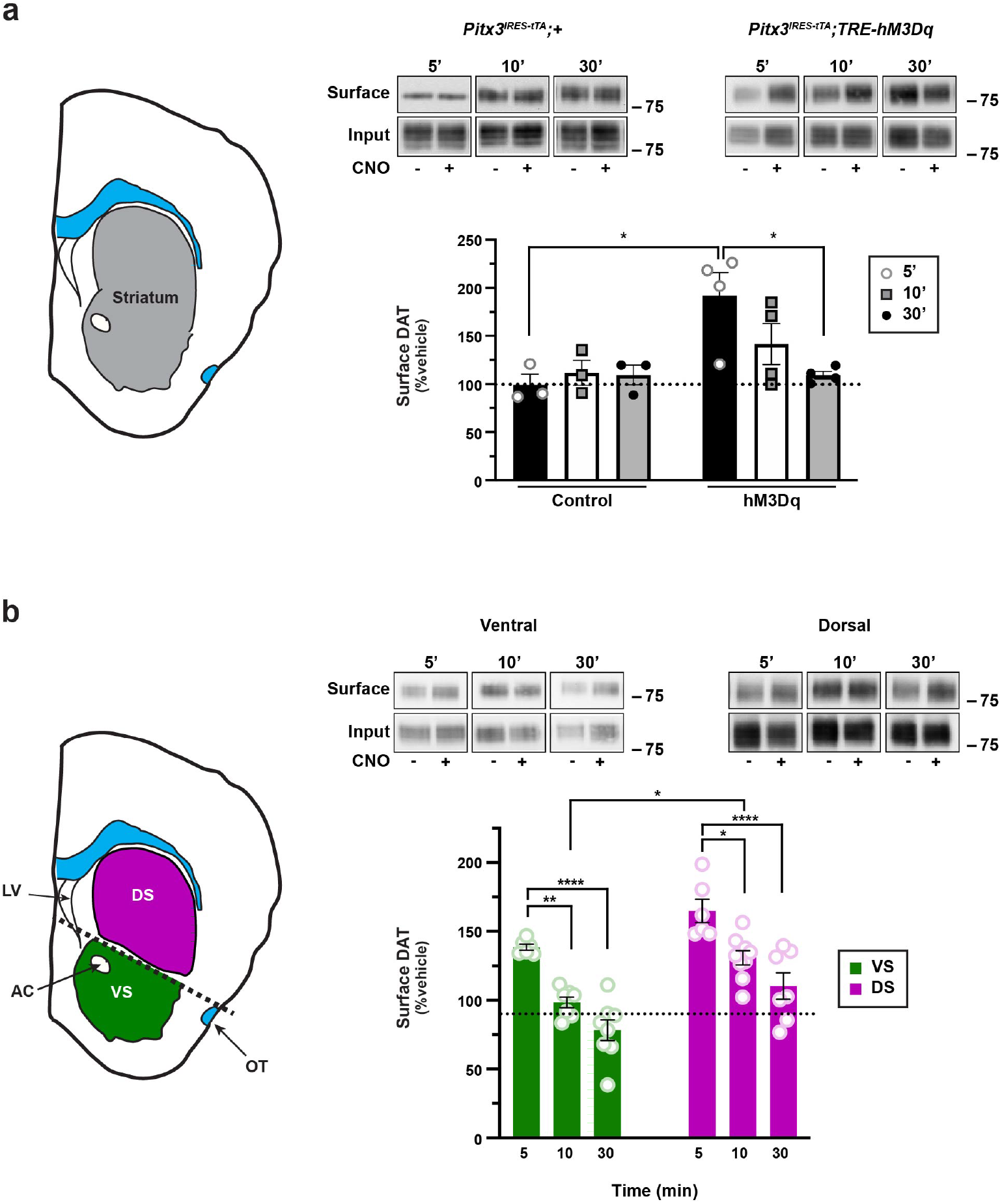
Gq-coupled DREADD activation drives region-specific, biphasic DAT trafficking. *Ex vivo striatal slice surface biotinylation.* Acute striatal slices were prepared from *Pitx3^IRES-tTA^;+* or *Pitx3^IRES-tTA^;TRE-hM3Dq* mice were treated ±500nM CNO for 5, 10 or 30 min, and DAT surface levels were measured by surface biotinylation as described in *Methods.* **(a)** *Total striatum.* Total striatal slices (*left)* were assessed for hM3Dq-mediated DAT trafficking. *Right, top:* Representative DAT immunoblots showing surface and input DAT bands. *Right bottom:* Mean DAT surface levels are presented as %vehicle-treated contralateral hemi-section ±S.E.M. Two-way ANOVA: Interaction: F_(2,15)_=3.82, *p=0.045; Genotype: F_(1,15)_=8.53, *p=0.011, Time: F_(2,15)_=2.24, p=0.24. *p<0.05, Tukey’s multiple comparisons test, n=3 (*Pitx3^IRES-tTA^*) and 4 (*Pitx3^IRES-tTA^;TRE-hM3Dq).* **(b)** *Sub-dissected striatum. Left:* Dorsal and ventral striata were sub-dissected prior to solubilizing, as described in *Methods. Right, top:* Representative DAT immunoblots showing surface and input DAT bands. *Right bottom:* Mean DAT surface levels, presented as %vehicle-treated contralateral hemi-section ±S.E.M. Two-way ANOVA: Interaction: F_(2,37)_=0.12, p=0.89; Time: F_(2,37)_=34.65, ****p<0.0001, Region: F_(1,37)_=30.05, ****p<0.0001). **p=0.003; *p=0.011, Tukey’s multiple comparisons test. *Ventral*: n=6 (5 min), 7 (10 min), and 8 (30 min); *Dorsal:* n=6 (5 min), 9 (10 min), and 8 (30 min).

### Biphasic Gq-stimulated DAT trafficking is mediated by DRD2 and PKC

Gq-stimulated, biphasic DAT trafficking was surprising in light of copious previous reports from both our laboratory^24, 25, 27, 29, 36^ and others^21, 30^ that direct PKC activation stimulates DAT internalization and decreases DAT surface expression. Moreover, Gq-coupled receptor activation in non-DAergic cells and midbrain decreases DAT surface levels in a PKC-dependent manner^30, 31^. Since our current studies tested Gq-stimulated DAT trafficking in DAergic terminals, we considered what factors, specific to DAergic terminals, might mediate biphasic DAT trafficking. Previous studies independently reported that 1) hM3Dq evokes DA release^37–39^, and 2) DA receptor 2 (DRD2) activation drives DAT insertion in synaptosomes^40–42^. Therefore, we hypothesized that 1) initial DAT membrane insertion may be due to hM3Dq-stimulated DA release and subsequent DRD2 autoreceptor (DRD2_auto_) activation, and 2) DAT subsequent return to baseline is mediated by PKC-stimulated DAT internalization, which may act as a retrieval mechanism following enhanced DAT cell surface delivery.

To test these hypotheses, we first asked whether DA release is required for hM3Dq-stimulated DAT insertion. To block DA release, we depleted vesicular DA stores with a single reserpine injection (5 mg/kg, I.P.) 16 hours prior to preparing striatal slices, which is sufficient to block evoked DA release^43, 44^. Vesicular DA depletion completely abolished hM3Dq-stimulated DAT insertion in response to a 5 min CNO treatment, in both VS and DS, as compared to slices from saline-injected mice (Fig. 2a). Interestingly, reserpine treatment also increased basal DAT surface expression in the DS, but not VS (Fig. S1a). These results demonstrate that vesicular DA release is required for rapid hM3Dq-stimulated DAT insertion.

**Figure 2.**
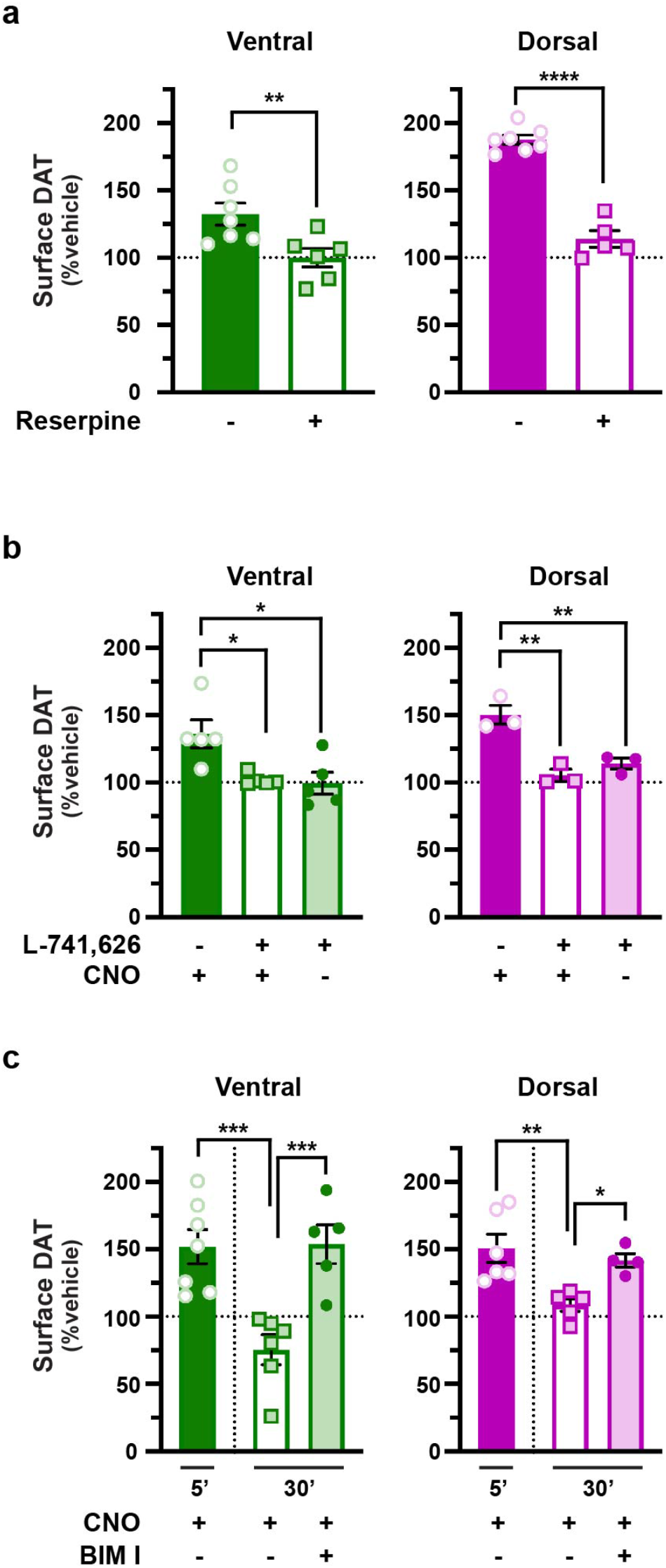
DA release and DRD2 activation are required for Gq-stimulated DAT insertion, whereas PKC activity is required for DAT retrieval. *Ex vivo striatal slice surface biotinylation.* Acute striatal slices prepared from *Pitx3^IRES-tTA^;TRE-hM3Dq* mice were treated with the indicated drugs for the indicated times. DAT surface levels were measured by slice biotinylation and VS and DS were isolated prior to tissue lysis as described in *Methods.* Mean DAT surface levels in ventral (*left)* and dorsal (*right)* striata are presented as %vehicle-treated contralateral hemi-section ±S.E.M. **(a)** *Reserpine treatment.* Mice were injected (I.P.) ±5.0 mg/kg reserpine 16 hrs prior to preparing slices, and 1.0µM reserpine or vehicle were included in the bath throughout the experiment. Slices were treated ±500nM CNO, 5 min, 37°C. *Ventral:* **p=0.007, one-tailed, unpaired Student’s t test, n=8 (saline) and 6 (reserpine). *Dorsal:* ****p<0.0001, one-tailed, unpaired Student’s t test, n=7 (saline) and 5 (reserpine). **(b)** *DRD2 antagonist pretreatment.* Striatal slices were pretreated ±DRD2 antagonist (L-741,626, 25nM, 15 min, 37°C), then treated ±500nM CNO (5 min, 37°C). DRD2 blockade abolished hM3Dq-stimulated DAT membrane insertion, but had no effect alone in either ventral (*left)* or dorsal (*right*) striata. *Ventral:* Kruskal-Wallis test, 8.82. *p<0.05, Dunn’s multiple comparisons test, n=5. *Dorsal:* One-way ANOVA, F_(2,6)_=20.17, **p=0.002. **p<0.01, Bonferroni’s multiple comparisons test, n=3. **(c)** *PKC inhibition.* Slices were pretreated ±500nM CNO (5 min, 37°C), followed by treatment ±1.0µM BIM I (25 min, 37°C), and DAT surface levels were measured by slice biotinylation as described in *Methods. Ventral striatum.* BIM I significantly blocked DAT retrieval following CNO-stimulated membrane insertion in both ventral and dorsal striata. *Ventral:* One-way ANOVA: F_(3,19)_=9.33, ***p=0.0005. ***p<0.002, Bonferroni’s multiple comparisons test, n=5-7. *Dorsal:* One-way ANOVA: F_(3,14)_=9.41, **p=0.001. *p=0.04, **p=0.004, Bonferroni’s multiple comparisons test, n=6-7.

We next asked whether DRD2 activation is required for hM3Dq-stimulated DAT insertion in VS and DS, and/or is sufficient to drive rapid DAT insertion. Slices were pretreated with the DRD2-specific antagonist L-741,626^45^ (25nM, 15 min, 37°C) and hM3Dq-stimulated DAT insertion was evoked with CNO (500nM, 5 min, 37°C). DRD2 inhibition completely abolished CNO-stimulated DAT insertion in both VS and DS as compared to either vehicle-treated slices, or slices treated with L-741,626 alone (Fig. 2b). Further, direct DRD2 activation with the DRD2 selective agonist sumanirole^46^ (170nM, 5 min, 37°C) was sufficient to drive rapid DAT membrane insertion in both VS and DS in striatal slices prepared from wildtype mice (Fig. S1b). Enhanced DAT surface levels were sustained in the DS out to 30 min, but rapidly returned to baseline by 10 minutes in the VS (Fig. S1b), revealing that DAT trafficking kinetics are also regionally distinct in response to direct DRD2 activation. Previous studies reported that DRD2-stimulated DAT insertion in striatal synaptosomes requires PKCβ activity^40^. To test whether hM3Dq-stimulated DAT insertion was similarly dependent upon PKCβ activity, we pretreated slices with the PKCβ-specific inhibitor ruboxistaurin^47, 48^ (50nM, 30 min, 37°C), prior to CNO-stimulated DAT insertion. Ruboxistaurin pretreatment completely abolished CNO-stimulated DAT insertion as compared to vehicle-pretreated slices (Fig. S1c), consistent with DRD2-mediated, PKCβ−dependent DAT membrane delivery. Taken together, these data demonstrate that both DA release and DRD2 activation are required for hM3Dq-stimulated DAT membrane insertion in VS and DS.

We next investigated whether DAT retrieval following hM3Dq-stimulated insertion was mediated by PKC-mediated DAT internalization. To test this, we first induced DAT insertion with CNO (500nM, 5 min), and then treated slices ± the PKC-specific inhibitor BIM I (1µM). Although CNO treatment alone drove biphasic DAT insertion and subsequent retrieval in control slices, PKC inhibition significantly abolished DAT return to baseline in both VS and DS, (Fig. 2c), consistent with the premise that DAT retrieval following hM3Dq-stimulated insertion is mediated by PKC-stimulated DAT internalization. BIM I treatment alone had no effect on DAT surface expression (Fig. S1d), suggesting that there is little tonic PKC-stimulated DAT internalization in *ex vivo* slices. Taken together these results clearly demonstrate that presynaptic Gq-coupled receptor activation drives biphasic DAT trafficking that is facilitated by Gq-stimulated DA release, DRD2-mediated DAT insertion, and PKC-stimulated DAT internalization.

### mGluR5 drives biphasic DAT trafficking that requires DRD2_auto_, Vps35, and Rit2

We next explored whether a native, presynaptic, Gq-coupled receptor similarly drives biphasic DAT trafficking. Group I metabotropic glutamate receptors (mGluR1 and 5) are Gq-coupled GPCRs that are highly expressed in striatum and midbrain DA neurons^49–51^. Moreover, a previous report found that mGluR5 activation functionally downregulated DAT in striatal synaptosomes^32^. To test whether Group I mGluRs regulate DAT surface expression, we treated *ex vivo* striatal slices from wildtype mice with the Group I mGluR agonist, DHPG (10µM). Similar to our results with hM3Dq, DHPG treatment drove rapid DAT membrane insertion at 5 min in both VS and DS (Fig. 3a, b). In VS, surface DAT was significantly retrieved by 10 min, similar to the hM3Dq-stimulated trafficking. However, in DS, DAT retrieval was considerably more rapid following DHPG-stimulated insertion than it was in response to hM3Dq activation, with a significant decrease to baseline by 10 min, and no further change by 30 min (Fig. 3b). To test whether DHPG-stimulated DAT insertion at 5 min was specifically mediated by mGluR5, mGluR1, or both, we pretreated slices with the mGluR5-specific antagonist MTEP^52^ (50nM, 15 min) prior to DHPG treatment. MTEP pretreatment completely abolished DHPG-stimulated DAT insertion in both VS and DS as compared to slices treated with DHPG alone (Fig. 3c, d), consistent with an mGluR5-, but not mGluR1-, mediated mechanism.

**Figure 3.**
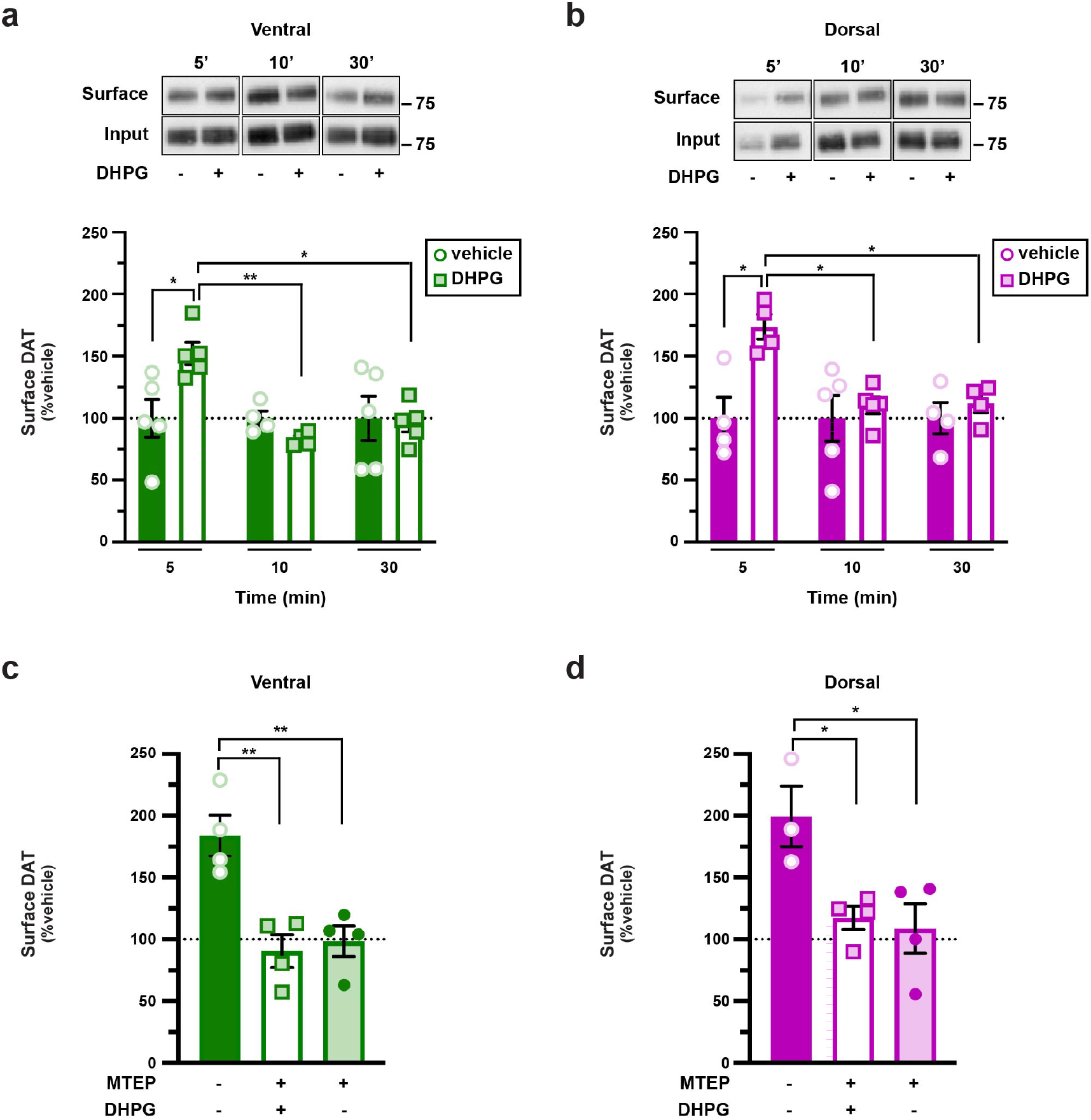
Striatal mGluR5 activation drives biphasic DAT trafficking. *Ex vivo striatal slice surface biotinylation.* Acute striatal slices prepared from male C57Bl/6J mice were treated with the indicated drugs for the indicated times. DAT surface levels were measured by slice biotinylation and dorsal and ventral striata were sub-dissected prior to solubilizing as described in *Methods.* Mean surface DAT levels are expressed as %vehicle-treated contralateral hemi-section ±SEM. **(a, b)** *DHPG treatment.* Slices were treated ±10µM DHPG for 5, 10 or 30 min. *Tops*: Representative immunoblots. *Bottoms:* Mean values are presented as. **(a)** *Ventral:* Two-way ANOVA: Interaction: F_(2,22)_=4.95, *p=0.02; Time: F_(2,22)_=4.95, *p=0.02, Drug: F_(1,22)_=2.24, p=0.29. *p<0.05, **p=0.006, Tukey’s multiple comparisons test, n=4-5x. **(b)** *Dorsal:* Two-way ANOVA: Interaction: F_(2,20)_=3.65, *p=0.04; Time: F_(2,20)_=3.65, *p=0.04, Drug: F_(1,20)_=8.80, **p=0.008. *p<0.05, Tukey’s multiple comparisons test, n=4-5. **(c, d)** *mGluR5 antagonist pretreatment.* Slices were pretreated ±MTEP (50nM, 15 min, 37°C), a selective mGlur5 antagonist, prior to stimulating DAT insertion with DHPG (10µM, 5 min, 37°C). MTEP pretreatment abolished DHPG-stimulated DAT membrane insertion in both ventral (c) and dorsal (d) striata. *Ventral:* One-way ANOVA: F_(2,9)_=13.42, **p=0.002; **p<0.01, Bonferroni’s multiple comparisons test, n=4. *Dorsal:* One-way ANOVA: F_(2,8)_=6.92, *p=0.02. *p<0.05, Bonferroni’s multiple comparisons test, n=3-4.

### Presynaptic mGluR5 drives DAT trafficking and influences DAT surface tone

mGluR5 is widely expressed in the striatum and is reportedly expressed in midbrain neurons. However, it is unknown whether mGluR5 is expressed in DAergic terminals; thus, it is unclear whether DHPG stimulates DAT trafficking directly via a presynaptic mGluR5, or indirectly via mGluR5s expressed elsewhere within the striatum. We previously leveraged the Tet-OFF approach to achieve AAV-mediated, conditional gene silencing in DA neurons^29, 35^. We demonstrated that AAV9 injection into *Pitx3^IRES-tTA^* mouse ventral tegmental area (VTA) transduced dopamine neurons in both VTA and substantia nigra pars compacta (SNc) and achieved robust gene silencing in SNc, but not in the neighboring non-DAergic SN reticulata, confirming both the viral spread and selectivity for DA neurons. To further assure that injecting our AAV9 vectors into VTA efficaciously transduced nigrostriatal projections into the DS, as well as mesolimbic projections to the VS, we assessed the striatal GFP reporter expression encoded in our AAV9 vectors. AAV9 particle injection into VTA resulted in widespread, robust GFP expression throughout both DS and VS, that co-localized with TH+ terminals (Fig. S2a).

To directly test whether mGluR5 regulation of DAT was cell-autonomous, we leveraged this approach to conditionally silence mGluR5 in DA neurons. *Pitx3^IRES-tTA^;mGluR5^fl/fl^*mouse VTA were bilaterally injected with AAV9-TRE-eGFP or -TRE-Cre, and we tested whether DHPG-stimulated DAT trafficking in VS and DS ex vivo slices requires presynaptic. AAV9-TRE-Cre significantly reduced midbrain mGluR5 mRNA expression as compared to eGFP-injected controls (Fig. 4a). While mGluR5 activation (10µM DHPG, 5 min, 37°C) drove DAT insertion in VS and DS slices from control mice (Fig. 4b, c), DAergic mGluR5 silencing completely abolished DHPG-stimulated DAT insertion in both VS and DS (Fig. 4b, c), consistent with the premise that mGluR5 is expressed presynaptically in DA terminals, where it stimulates DAT trafficking via a cell-autonomous mechanism. We further hypothesized that DAT surface tone in the striatum is a balance between DRD2-stimulated DAT insertion and mGluR5-mediated retrieval, and therefore predicted that mGluR5 silencing in DA neurons would lead to enhanced basal DAT surface expression. To test this possibility, we measured basal DAT surface levels following conditional mGluR5 silencing. DAergic mGluR5 silencing significantly increased baseline DAT surface levels in both VS and DS, as compared to slices from control mice (Fig. 4d), consistent with the ability of DAT to undergo DRD2-stimulated insertion, but not Gq-stimulated retrieval. These data support the hypothesis that mGluR5-stimulated DAT retrieval significantly impacts striatal DAT surface tone.

**Figure 4.**
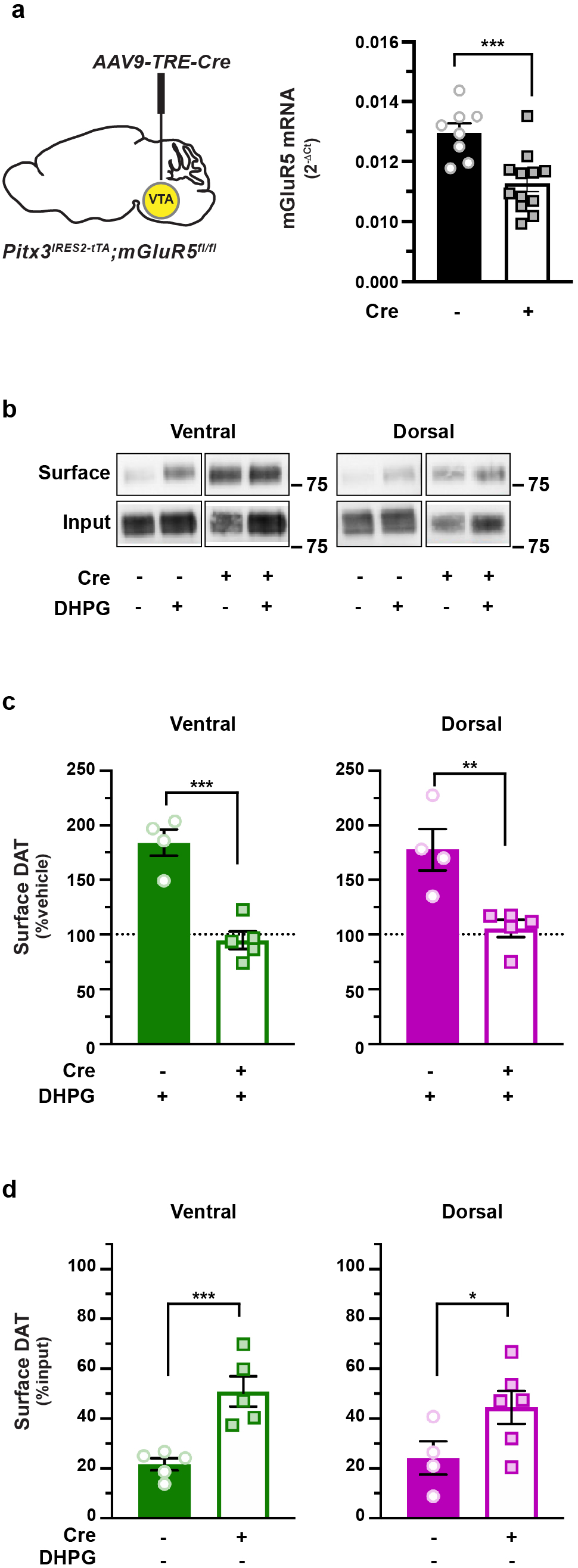
mGluR5-mediated DAT trafficking is mediated presynaptically and impacts basal DAT surface expression. **(a)** *Conditional mGluR5 silencing in DA neurons. (Left) Pitx3^IRES-tTA^;mGluR5^fl/fl^*mouse VTA were bilaterally injected with either AAV9-TRE-eGFP (n=7) or AAV9-TRE-Cre (n=10). *(Right)* Midbrain mGluR5 mRNA levels were measured by RT-qPCR from tissue punches obtained 4-5 weeks post-injection. ***p=0.0005, one-tailed, unpaired Student’s *t* test, n=8 (Cre) and 12 (eGFP). **(b-d)** *Ex vivo striatal slice surface biotinylation. Pitx3^IRES-tTA^;mGluR5^fl/fl^* mouse VTA were bilaterally injected with either AAV9-TRE-eGFP (n=4) or AAV9-TRE-Cre (n=5). Acute striatal slices were prepared from the indicated mice, treated ±10µM DHPG (5 min, 37°C), and DAT surface levels were measured by slice biotinylation as described in *Methods.* Slices were sub-dissected into ventral and dorsal striatal subregions prior to solubilizing. **(b)** *Representative immunoblots:* Surface and input DAT bands are presented for each of the indicated treatment conditions. **(c)** *DHPG-stimulated DAT membrane insertion:* Mean DAT surface levels are presented as %vehicle-treated contralateral hemi-section. DAergic mGluR5 silencing significantly abolished DHPG-stimulated DAT insertion in both ventral (***p=0.0002), and dorsal (**p=0.003) striata, one-tailed, unpaired Student’s t test. **(d)** *Basal DAT surface expression:* Mean surface DAT values are presented as %input surface levels. DAergic mGluR5 silencing significantly increased basal DAT surface expression in ventral (***p=0.001) dorsal (*p=0.035) striata, one-tailed, unpaired Student’s t test, n= 5 (VS) and 4-6 (DS).

### Biphasic mGluR5-stimulated DAT trafficking requires DRD2auto, retromer, and Rit2

While hM3Dq-stimulated DAT insertion and retrieval are DRD2- and PKC-dependent, respectively, it is unknown whether mGluR5-stimulated DAT trafficking was likewise dependent upon these mechanisms. It is further unknown whether DRD2-stimulated DAT insertion is mediated cell autonomously by the presynaptic DRD2_auto_, or cell non-autonomously by DRD2 activation elsewhere within the striatum, where DRD2s are widely expressed. To discriminate between these possibilities, we aimed to conditionally excise DRD2_auto_ in a *DRD2^fl/fl^* mouse, and test whether DRD2_auto_ was required for mGluR5-stimulated DAT insertion. Given previous reports that Cre-mediated DRD2_auto_ excision at germline significantly enhances DA release^53^, we again opted to leverage the Tet-OFF approach to virally excise DRD2_auto_ post-development in *Pitx3^IRES-tTA^;DRD2^fl/fl^* mice, and measure DAT trafficking in response to mGluR5 activation. AAV9-TRE-Cre injection into *Pitx3^IRES-tTA^;DRD2^fl/fl^* mouse VTA significantly decreased midbrain *DRD2* mRNA expression, consistent with DRD2_auto_ excision (Fig. 5a), but did not significantly alter basal DAT surface expression in either DS or VS (Fig. S2b). In VS and DS, DRD2_auto_ silencing completely abolished DHPG-stimulated DAT insertion (Fig. 5b). Thus, DRD2_auto_ is specifically required for mGluR5-stimulated DAT insertion.

**Figure 5.**
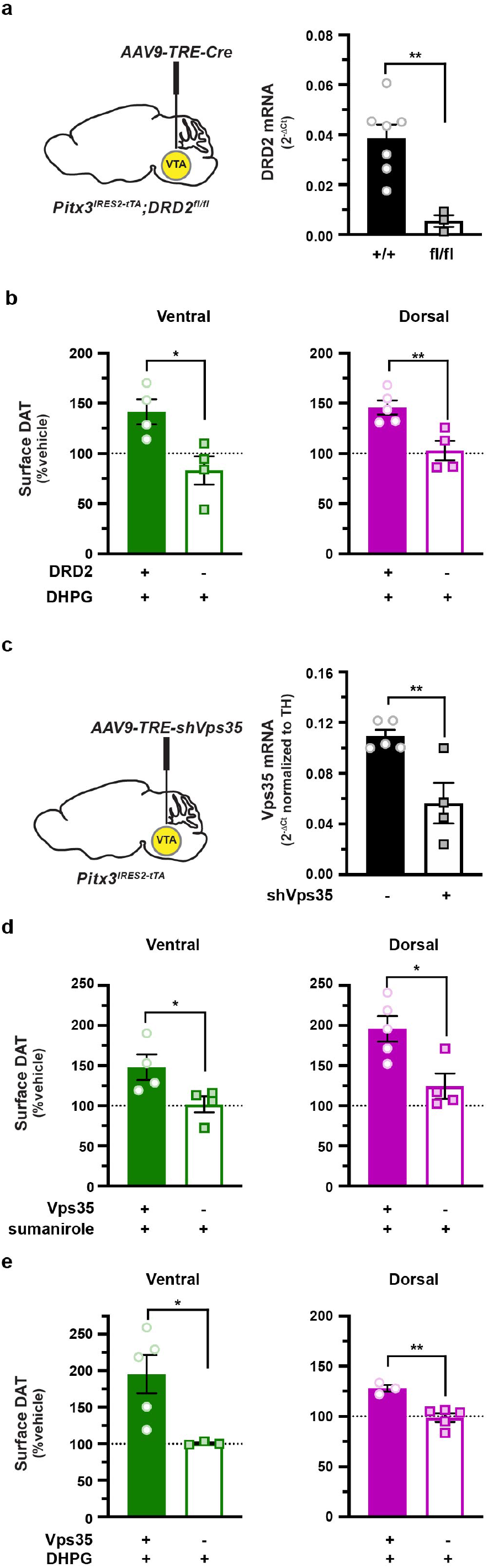
mGluR5-mediated DAT insertion requires DRD2_auto_ and intact retromer. **(a)** *Virally-induced DRD2 autoreceptor silencing. (Left) Pitx3^IRES-tTA^;DRD2^fl/fl^* mouse VTA were bilaterally injected with either AAV9-TRE-eGFP or AAV9-TRE-Cre. *(Right)* Midbrain *DRD2* mRNA levels were measured by RT-qPCR from tissue punches obtained 4-5 weeks post-injection. **p=0.002, one-tailed, unpaired Student’s *t* test, n=3-7. **(b)** *Ex vivo striatal slice surface biotinylation.* Acute striatal slices prepared from the indicated mice were treated ±10µM DHPG (5 min, 37*C), and DAT surface levels were measured by slice biotinylation as described in *Methods.* Surface DAT is expressed as a percentage of the contralateral vehicle-treated hemi-section ±SEM. DRD2 autoreceptor silencing abolished mGluR5-stimulated DAT membrane delivery in both ventral (*p=0.01) and dorsal (**p=0.004) striata, one-tailed, unpaired Student’s t test, n=4 (ventral) and 4-5 (dorsal). **(c-e)** *Conditional Vps35 silencing in DA neurons.* **(c,** *Left) Pitx3^IRES-tTA^*mouse VTA were bilaterally injected with either AAV9-TRE-eGFP or AAV9-TRE-shVps35. (**c**, *Right)* Midbrain Vps35 mRNA levels were measured by RT-qPCR from tissue punches obtained 4-5 weeks post-injection, and were normalized to TH mRNA levels to account for varied DA neuron enrichment amongst tissue punches. **p=0.005, one-tailed, unpaired Student’s t test, n=4-5. **(d, e)** *Ex vivo striatal slice surface biotinylation.* Acute striatal slices were prepared from the indicated mice, were treated with the indicated drugs (5 min, 37°C), and DAT surface levels were measured by slice biotinylation as described in *Methods.* Surface DAT is expressed as a percentage of the contralateral vehicle-treated hemi-section ±SEM. (**d)** *DRD2-stimulated DAT membrane delivery*: Slices were treated ±170nM sumanirole (5 min, 37°C). Vps35 silencing abolished DRD2-stimulated DAT membrane delivery in both ventral (*p=0.049) and dorsal (*p=0.02) striata, one-tailed, unpaired Student’s t test, n=4-5. (**e)** *mGluR5-stimulated DAT membrane delivery.* Slices were treated ±10µM DHPG (5 min, 37°C). Vps35 silencing abolished mGluR5-stimulated DAT membrane delivery in both ventral (*p=0.03) and dorsal (**p=0.003) striata, one-tailed, unpaired Student’s t test, n=3-5.

We next investigated the molecular mechanisms required for DRD2-and mGluR5-stimulated DAT insertion. We previously reported that internalized DAT targets to retromer/Vps35^+^ endosomes, and that intact retromer complex is required for DAT delivery to the plasma membrane in dopaminergic neuroblastoma cells lines^28^. We therefore hypothesized that retromer may likewise be required for both DRD2- and mGluR5-stimulated DAT surface delivery in DAergic terminals. To test this possibility, we conditionally silenced Vps35 in DA neurons via Tet-OFF, AAV-mediated delivery of a validated, mouse-directed Vps35 shRNA^54^, and tested the ability of either sumanirole or DHPG to induce DAT insertion in either DS or VS DAergic terminals. AAV9-TRE-shVps35 significantly diminished *Vps35* mRNA expression in *Pitx3^IRES-tTA^*mouse midbrain, as compared to control-injected mice (Fig. 5c), but did not significantly impact basal DAT surface expression in either DS or VS (Fig. S2c). DRD2- (Fig. 5d) and mGluR5-stimulated (Fig. 5e) DAT insertion in both VS and DS were significantly attenuated following DAergic Vps35 knockdown, as compared to slices from control-injected mice, demonstrating that retromer is required for both DRD2 and mGluR5-stimulated DAT insertion in DAergic terminals.

We previously reported that the neuronal GTPase Rit2 is required for phorbol ester-stimulated DAT endocytosis in cell lines and striatal slices^27, 29^. However, it is unknown whether Rit2 is required for PKC-mediated DAT retrieval following mGluR5-stimulated DAT surface delivery. We therefore leveraged our previously validated approach to conditionally silence DAergic Rit2 in *Pitx3^IRES-tTA^* mice^29, 35^ and tested whether RIt2 was required for either mGluR5-stimulated DAT insertion or retrieval. As previously reported, AAV9-TRE-shRit2 significantly reduced midbrain Rit2 mRNA (Fig. 6a). Rit2 is not known to be requisite for DAT insertion, therefore, we predicted that, in the absence of Rit2, mGluR5 activation would still deliver DAT to the plasma membrane, but that it would be unable to be retrieved. Indeed, DAergic Rit2 knockdown had no significant effect on mGluR5-stimulated DAT insertion as compared to slices from control-injected mice, in either VS (Fig. 6b) or DS (Fig. 6c). However, DAergic Rit2 silencing significantly blocked DAT retrieval and return to baseline in both VS and DS (Fig. 6b, c), indicating that DAergic Rit2 is required for PKC-mediated DAT retrieval in both striatal subregions.

**Figure 6.**
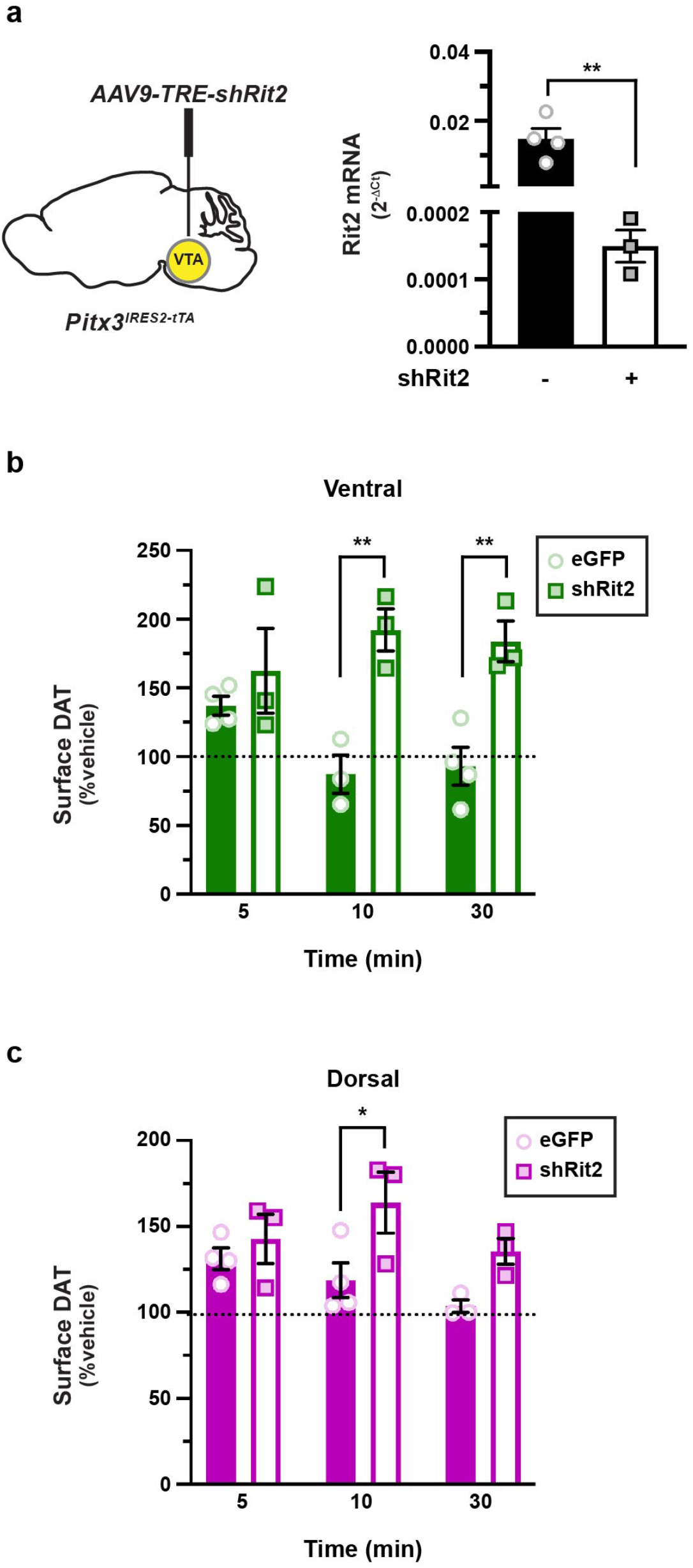
Rit2 is required for mGluR5-mediated DAT retrieval, but not insertion. **(a)** *Conditional Rit2 silencing in DA neurons. (Left) Pitx3^IRES-tTA^* mouse VTA were bilaterally injected with either AAV9-TRE-eGFP or AAV9-TRE-Cre. *(Right)* Midbrain *Rit2* mRNA levels were measured by RT-qPCR from tissue punches obtained 4-5 weeks post-injection. **p=0.005, one-tailed, unpaired Student’s *t* test, n=3-4. **(b, c)** *Ex vivo striatal slice surface biotinylation.* Acute striatal slices were prepared from the indicated mice, treated ±10µM DHPG for the indicated times (37°C), and DAT surface levels were measured by slice biotinylation as described in *Methods.* Slices were sub-dissected into ventral **(b)** and dorsal **(c)** striatal subregions prior to solubilizing. Surface DAT is expressed as a percentage of the contralateral vehicle-treated hemi-section ±SEM. **(b)** *Ventral striatum:* Two-way ANOVA: Interaction: F_(2,14)_=3.35, p=0.065; Time: F_(2,14)_=0.29, p=0.75, Virus: F_(1,14)_=30.00, ****p<0.0001). Rit2 silencing significantly abolished DAT retrieval at 10 min (**p=0.002) and 30 min (**p=0.004), Sidak’s multiple comparisons test, n=3-4. **(c)** *Dorsal striatum:* Two-way ANOVA: Interaction: F_(2,14)_=1.30, p=0.30; Time: F_(2,14)_=2.19, p=0.15, Virus: F_(1,14)_=11.44, **p=0.004. Rit2 silencing significantly abolished DAT retrieval at 10 min (*p=0.02), Sidak’s multiple comparisons test, n=3-4.

### DRD2-stimulated DAT insertion and presynaptic mGluR5 expression impact striatal DA dynamics

Despite decades-long research that demonstrate regulated DAT trafficking, it remains unknown whether or not DAT trafficking impacts DAergic signaling. To test this possibility, we leveraged fast-scan cyclic voltammetry (FSCV) in *ex vivo* dorsal striatal slices to ask whether 1) DRD2-mediated DAT insertion, or 2) mGluR5 modulation of DAT basal surface levels impacted DA release and/or clearance. To this end, we conditionally excised mGluR5 selectively from DA neurons in *Pitx3^IRES-tTA^;mGluR5^fl/fl^* mice injected with either AAV9-TRE-eGFP (control) or AAV9-TRE-Cre. We first probed how DRD2 activation impacts DA clearance in slices from control (eGFP) mice (Fig. 7a-d). We predicted that evoked DA release would drive DRD2-stimulated DAT insertion and would decrease DA clearance times in slices from control mice. Thus, we compared DA transients evoked in ACSF alone, to those evoked in the presence of the DRD2-specific antagonist L-741,626 (Fig. 7 a-d; see also Table I). Electrically evoked DA transients in the presence of L-741,626 (25nM) had an average amplitude 655.6 ±110.6nM (Fig. 7c), and those recorded in ACSF alone were significantly smaller (361.1 ±105.5nM, Fig. 7c), consistent with DRD2-mediated TH inhibition. DA clearance, measured as the exponential decay tau, was 0.44±0.06 sec in in the presence of L-741,626 (Fig. 7d). However, the decay tau was significantly shortened when recorded in ACSF alone (0.27 ±0.02 sec; Fig. 7d), consistent with our prediction that DRD2-stimulated DAT insertion would increase clearance rates.

**Figure 7.**
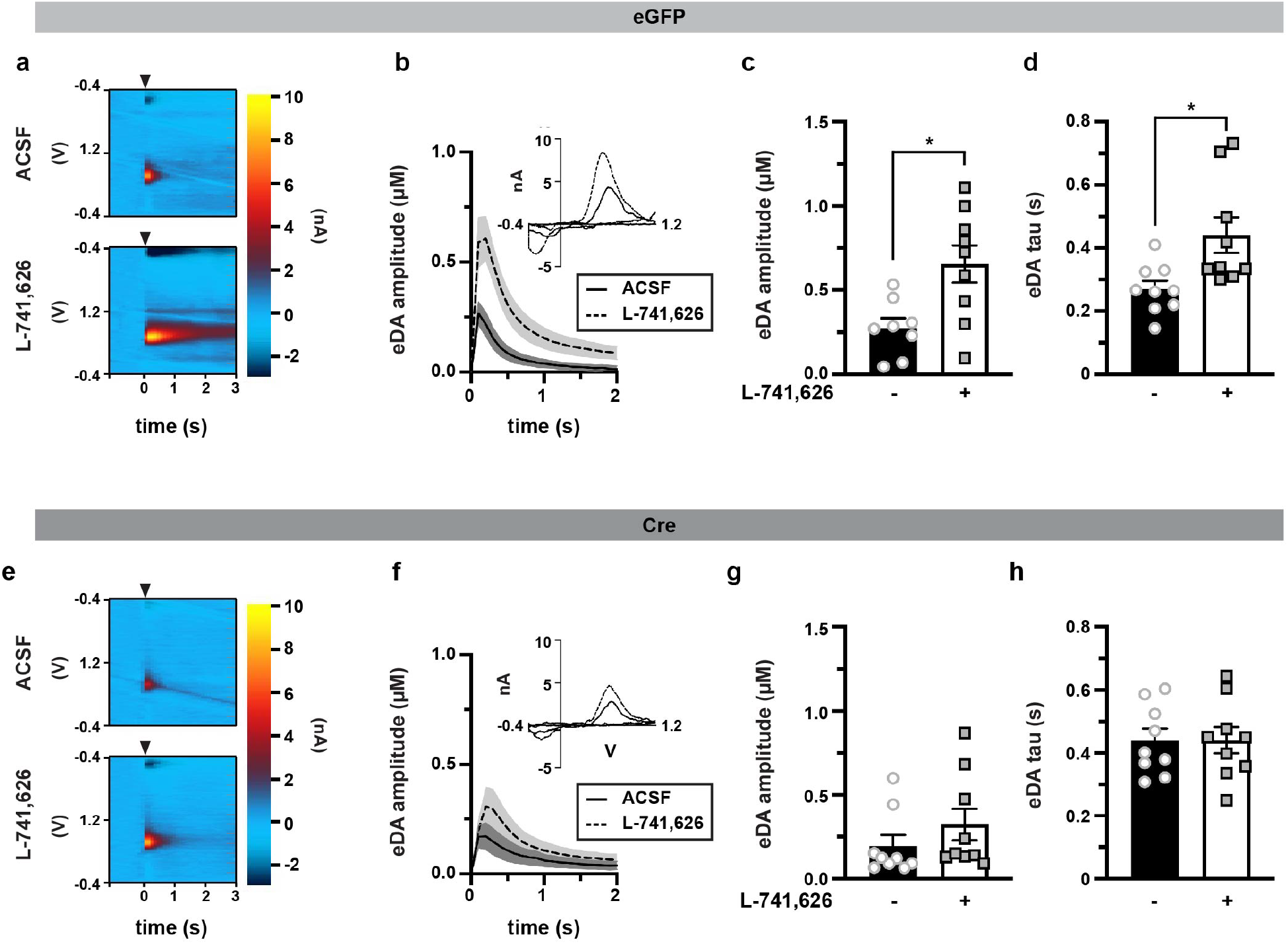
DRD2-stimulated DAT insertion and mGluR5-mediated DAT retrieval impact DA release and clearance in dorsal striatum. *Ex vivo fast-scan cyclic voltammetry: Pitx3^IRES-tTA^;mGluR5^fl/f^* mouse VTA were bilaterally injected with either AAV9-TRE-eGFP (n=7-9) or AAV9-TRE-Cre (n=9). Acute striatal slices were prepared from the indicated mice, and electrically evoked DA transients were measured in dorsal striatum by FSCV as described in *Methods.* **(a-d)** *eGFP:* **(a)** *Representative voltammograms:* Voltammograms displaying evoked current over voltage cycles and time, in slices from eGFP-injected in the presence of ASCF alone (*top)* or supplemented with 25nM L-741,626 (*bottom)*. Arrowheads indicated delivery of single, squared wave pulse. **(b)** *Dopamine transients:* Evoked DA transients in slices from eGFP-injected mice, treated ±L-741,626 (25nM). Mean traces are presented with S.E.M. indicated by shaded areas. **(c)** *Mean amplitudes:* Mean amplitudes are presented in µM ±S.E.M. *Significantly greater than in ACSF alone, p=0.02. **(d)** Mean decay tau, presented in sec ±S.E.M. *Significantly longer clearance time than in ACSF alone, p=0.03. **(e-h)** *Cre: Representative voltammograms:* Voltammograms displaying evoked current over voltage cycles and time, in slices from eGFP-injected in the presence of ASCF alone (*top)* or supplemented with 25nM L-741,626 (*bottom)*. Arrowheads indicated delivery of single, squared wave pulse. **(b)** *Dopamine transients:* Evoked DA transients in slices from Cre-injected mice, treated ±L-741,626 (25nM). Average traces are presented with S.E.M. indicated by shaded areas. **(c)** *Mean amplitudes:* Mean amplitudes are presented in µM ±S.E.M. DRD2 inhibition did not significantly alter DA release inhibition, p=0.71. **(d)** Mean decay tau, presented in sec ±S.E.M. DRD2 inhibition had no significant effect on DA clearance, p>0.999. See Table I for all descriptive statistical analyses.

**Table I.** Effect of DRD2 inhibition and DAergic mGluR5 silencing on evoked DA transients

| **L-741,626** | **(-)** | **(+)** |
| --- | --- | --- |
| **Amplitude** |  |  |
| *eGFP* | 274.1 ±59.3 | *655.6 ±110.6 |
| *Cre* | 194.9 ±64.0 | †322.9 ± 95.2 |
| **Tau (sec)** |  |  |
| *eGFP* | 0.27 ±0.26 | *0.44 ±0.06 |
| *Cre* | †0.44 ±0.04 | 0.44 ±0.04 |

*Amplitude*: Two-way ANOVA: Interaction: $F_{(1,31)}=2.15$ , $p=0.15$ ; Virus: $F_{(1,31)}=5.67$ , $p=0.02$ , Drug: $F_{(1,31)}=8.67$ , $p=0.006$ . Tukey's multiple comparison test, $n=8-9$ .
\*Significantly increased as compared to $eGFP_{(-L741,626)}$ slices, $p=0.02$ .
†Significantly decreased as compared to $eGFP_{(+L741,626)}$ slices, $p=0.04$ .
*Tau*: Two-way ANOVA: Interaction: $F_{(1,32)}=4.20$ , $p=0.048$ ; Virus: $F_{(1,32)}=4.15$ ; $p=0.05$ , Drug: $F_{(1,32)}=4.21$ , $p=0.048$ . Tukey's multiple comparison test, $n=9$ .
\*Significantly increased as compared to $eGFP_{(-L741,626)}$ slices, $p=0.03$ .
†Significantly increased as compared to $eGFP_{(-L741,626)}$ slices, $p=0.03$ .

We further tested whether the enhanced DAT surface expression that occurred in response to DAergic mGluR5 silencing (as shown in Figure 4) impacted DA release and/or clearance. We predicted that since DAergic mGluR5 silencing increased DAT surface levels, DA clearance rates would be significantly faster than in slices from control mice. We additionally predicted that DRD2 activation would either further increase DA clearance, or alternatively would not impact clearance due to any potential ceiling effects on DAT membrane insertion. Results are shown in Figure 7e-h, and also Table I. To our surprise, following presynaptic mGluR5 silencing, DRD2 inhibition had no effect on the average DA transient amplitude as compared to those acquired in ACSF alone (Fig. 7g), and DA amplitudes measured in ASCF alone were not significantly different in slices from GFP-vs. Cre-injected mice (Table I). Similarly, DRD2 inhibition had no significant effect on decay tau values following DAergic mGluR5 silencing (Fig. 7h, p>0.999), Moreover, despite increased DAT surface expression in quiescent slices, decay tau values in Cre-injected mice were significantly longer than those in eGFP-injected mice (Table I, p=0.03), rather than shorter. These unexpected results could not be explained by alterations in DAergic tone, as DAergic mGluR5 silencing did not significantly affect either DAT or TH protein levels in either VS (Fig. S3a) or DS (Fig. S3b). Moreover, DAergic mGluR5 silencing did not affect pSer40-TH in either striatal region (Fig. S3a,b), suggesting that TH activity can be properly regulated in the absence of mGluR5.

### mGluR5-stimulated DAT trafficking is required for motor learning

In light of our new understanding that DAT undergoes biphasic surface trafficking in DAergic terminals, we next aimed to test whether disrupting mGluR5-mediated, biphasic DAT trafficking impacted DA-dependent motor behaviors. *Pitx3^IRES-tTA^;mGluR5^fl/fl^* mouse VTA were injected with either AAV9-TRE-Cre or -eGFP and mice were assessed across a battery of locomotor assays. mGluR5 loss from DA neurons did not significantly affect baseline horizontal locomotion (Fig. S4a, b), nor did it affect vertical (Fig. S4c) or fine (Fig. S4d) movements. However, DAergic mGluR5 silencing significantly perturbed motor learning on the accelerating rotarod as compared to control-injected mice (Fig. 8a), and also significantly decreased performance on the fixed speed rotarod (Fig. 8b), but not on the challenge balance beam, measured as both number of foot faults (Fig. 8c) and average traversal time (Fig. 8d). Interestingly, although average traversal time over 3 trials was unaffected by DAergic mGluR5 silencing, we observed that control mice significantly decreased their traversal time between the 1^st^ and 2^nd^ trials (Fig. 8e, *left*), whereas DAergic mGluR5 excision mice failed to improve their traversal time between the 1^st^ and 2^nd^ trials (Fig. 8e, *right*). This suggests that although DAergic mGluR5 excision does not affect coordination per se, it may impact other unknown aspects of balance beam performance, such as learning and/or habituation on test day. We further tested whether decreased rotarod performance was due to either muscle weakness/fatigue or gait disturbance, using the grip strength assay and gait assessment assays, respectively. DAergic mGluR5 silencing caused a modest, but significant, decrease in grip strength (Fig. S4e), but had no impact on gait as compared to control-injected mice, as measured by stride length, stride width, and toe spread, in both forelimbs and hindlimbs (Fig. S5). Thus, although grip strength was modestly affected, DAergic mGluR5 silencing overall had no impact on locomotion and/or gait, and therefore rotarod performance likely reflects deficits in motor learning.

**Figure 8.**
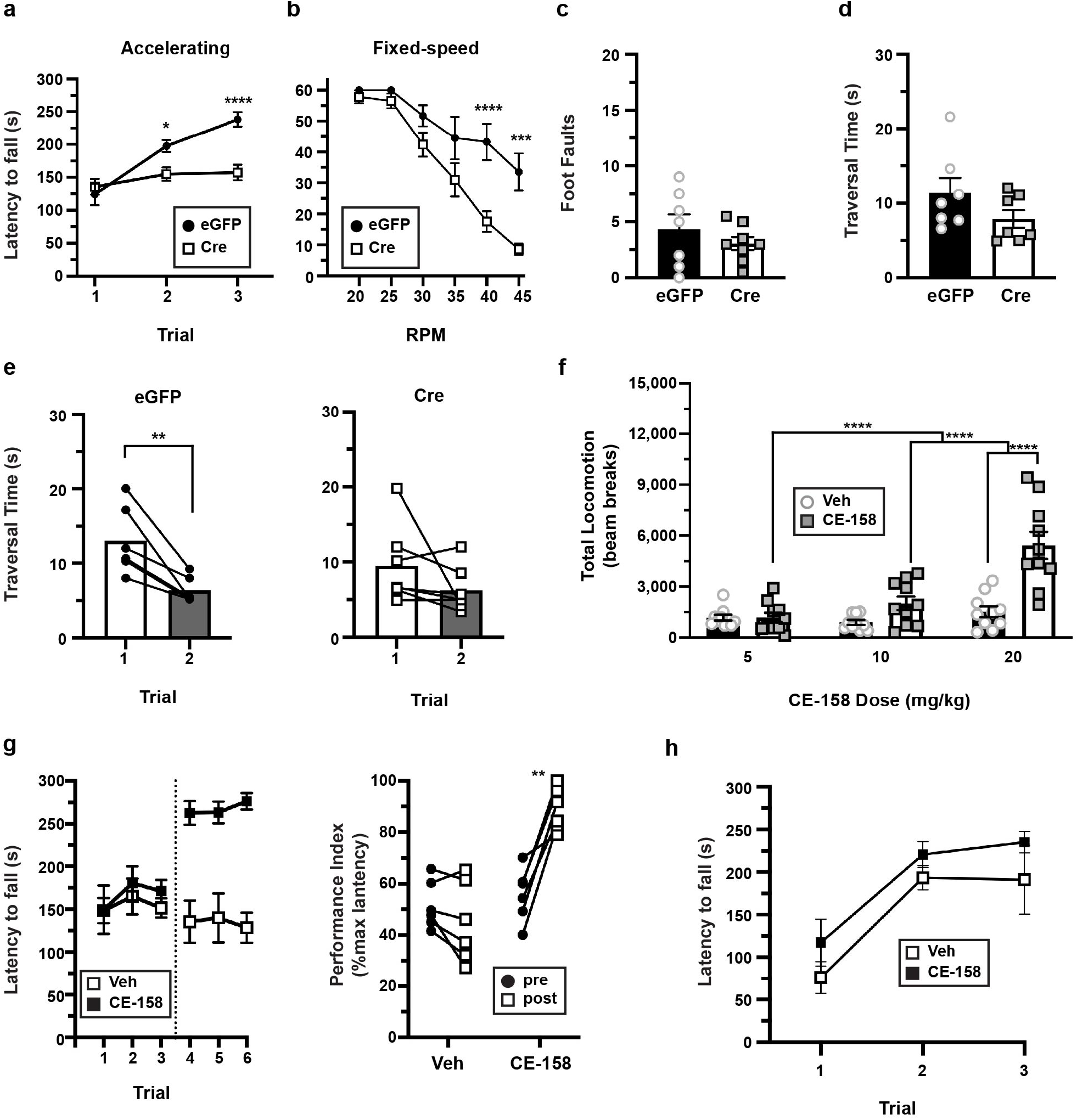
DAergic mGluR5 is required for motor learning in a DAT-dependent manner. *Pitx3^IRES-tTA^;mGluR5^fl/fl^* mouse VTA were bilaterally injected with either AAV9-TRE-eGFP (n=7-9) or AAV9-TRE-Cre (n=9) and were assessed by the indicated behavioral assays **(a)** *Accelerating Rotarod:* Mice were assessed over 3 consecutive trials as described in *Methods.* Two-way repeated measures ANOVA: Trial x Virus: F_(2,30)_=17.04, ****p<0.0001; Trial: F_(2,30)_=38.22, ****p<0.0001, Virus: F_(1,15)_=6.65, *p=0.02, Subject: F_(15,30)_=5.02, ****p<0.0001). mGluR5 silencing in DA neurons significantly dampened performance on Trials 2 (*p=0.049) and 3 (****p<0.0001), Bonferroni’s multiple comparisons test. **(b)** *Fixed-speed Rotarod:* Mice were assessed over the indicated consecutive speeds as described in *Methods.* Two-way repeated measures ANOVA: Speed x Virus: F_(5,75)_=5.06, ****p<0.0005; Speed: F_(5,75)_=44.91, ****p<0.0001, Virus: F_(1,15)_=12.23, **p=0.003, Subject: F_(15,15)_=4.09, ****p<0.0001. mGluR5 silencing in DA neurons significantly dampened performance at 40 rpm (****p<0.0001) and 45 (***p=0.0002), Bonferroni’s multiple comparisons test. **(c-e)** *Challenge balance beam.* Mean foot fault numbers **(c)** and beam traversal time (seconds) **(d)** are presented. mGluR5 silencing in DA neurons did not significantly affect either foot fault number (p=0.78) or mean traversal times (p=0.15), two-tailed, unpaired Student’s t test. **(e)** *Traversal time improvement.* eGFP (*control, left*) mouse traversal times significantly improved between trials 1 and 2 (*p=0.02), whereas mice injected with Cre (*right)* failed to improve their performance (p=0.77), two-tailed, paired Student’s t test. **(f)** *Dose-dependent effect of CE-158 on horizontal locomotion in wildtype mice.* Wildtype mice were habituated to PACs for 45 min, were then injected (I.P.) with either vehicle (on day 1) or the indicated CE-158 dose (on day 2), and horizontal locomotion measured for 90 min. Cumulative locomotion is presented as total post-injection beam breaks. Two-way ANOVA: Dose x Drug: F_(2,54)_=12.01, ****p<0.0001; Dose: F_(2,54)_=18.70, ****p<0.0001, Drug: F_(1,54)_=25.36, ****p<0.0001. 20mg/kg CE-158 significantly increased locomotion as compared to vehicle (****p<0.0001), 10mg/kg (****p<0.0001) and 5mg/kg (****p<0.0001), Bonferroni’s multiple comparisons test, n=9-10. **(g)** *Accelerating Rotarod rescue studies with CE-158:* Mice with DAergic mGluR5 silencing were assessed over 3 trials, injected I.P. with either saline or CE-158 (10mg/kg), and re-assessed over 3 trials 15 min post-injection. *Left:* Raw Rotarod results. *Right:* Rotarod Performance Indices, expressed as %maximal latency to fall. Mice injected with CE-158 performed significantly better post-injection as compared to pre-injection (**p=0.0014) whereas saline-injected mice did not perform significantly better (p=0.11). Two-tailed paired t test, n=6. **(h)** *Effect of CE-158 on wildtype mouse performance on accelerating Rotarod.* Wildtype mice were injected with either saline or CE-158 (10mg/kg) and assessed on the accelerating Rotarod. CE-158 had no significant effect on mouse learning over 3 trials. Two-way ANOVA: Trial x Drug: F_(2,22)_=0.085, p=0.92; Trial: F_(2,22)_=18.17, p<0.0001, Drug: F_(1,11)_=3.136, p=0.11, n=6 (veh) or 7 (CE-158).

Motor learning requires DA^55^ and the failure to perform on the accelerating rotarod following DAergic mGluR5 silencing led us to hypothesize that mGluR5-mediated DAT retrieval may be critical to boost extracellular DA levels and facilitate the learning process. Failure to retrieve DAT from the cell surface would be predicted to clear DA too quickly from the extracellular space, and not provide sufficient DA to facilitate the learning process. Indeed, enhanced basal surface DAT following DAergic mGluR5 silencing (Fig. 4d) is consistent with this hypothesis. Alternatively, mGluR5 activation in DAergic terminals could be playing a role in motor learning that is independent of DAT trafficking. To discriminate between these possibilities, we aimed to selectively block DAT activity with a sub-threshold dose of a DAT-specific inhibitor, which we predicted would rescue motor learning if enhanced DAT surface expression and function was solely responsible for rotarod deficits in mice with DAergic mGluR5 silencing. The majority of DAT inhibitors are also equipotent norepinephrine transporter (NET) inhibitors. However, a novel compound, (S,S)-CE-158, was recently reported that is highly selective for DAT over NET, permeates the blood-brain barrier, and increases extracellular DA levels in striatum following I.P. injection in rats^56^. We first tested what CE-158 doses are required to block DAT in mice *in vivo*, as assessed by its ability to drive hyperlocomotion, a hallmark of enhanced extracellular DA in rodent models. Wildtype mice were injected (I.P.) with either 5, 10, or 20mg/kg CE-158 and horizontal locomotion was measured. Only the 20mg/kg CE-158 dose resulted in significantly increased locomotion as compared either vehicle, 5mg/kg, or 10mg/kg injections (Fig. 8f, S6). Given the dose-response of CE-158, we next tested whether a subthreshold CE-158 dose (10 mg/kg) could rescue motor learning in mice with DAergic mGluR5 silencing. Mice were initially assessed on the rotarod, were injected with either vehicle or CE-158 (10 mg/kg, I.P.), and reassessed 15 min post-injection. CE-158 significantly improved mouse performance on the accelerating rotarod, whereas injection with vehicle had no significant effect on performance (Fig. 8g). Importantly, 10 mg/kg CE-158 (I.P.) had no impact on wildtype mouse performance on the accelerating rotarod (Fig. 8h). Thus, CE-158 does not impact motor learning per se, but selectively rescues motor learning following DAergic mGluR5 silencing, consistent with a potential role for DAT internalization dysfunction in motor learning.

## Discussion

Biphasic DAT trafficking has the potential to mold the DA signal during the ebb and flow of plastic events throughout the striatum. Our results point to a previously unknown role for glutamatergic signaling in the striatum to shape the temporal profile of DA signaling via DAT trafficking. Our emerging model is depicted in Figure 9. We hypothesize that under basal conditions, where DA neurons fire tonically, DAT is robustly delivered to the plasma membrane via DRD2_auto_ activation, in a PKCβ and retromer-dependent manner. During periods where DA demand is higher, such as during motor learning, glutamate release from striatal glutamatergic afferents activates mGluR5 on DAergic terminals, leading to DAT retrieval, decreased DAT surface expression, and an enhanced extracellular DA signal. This simple model does not take into account the plethora of presynaptic GPCRs, including Group II and III mGluRs, that also likely influence DAT surface expression. However, given DAT’s ability to rapidly traffic, it appears certain that presynaptic receptor signals continually integrate to orchestrate DAT surface levels and, ultimately, the extracellular DA signal.

**Figure 9.**
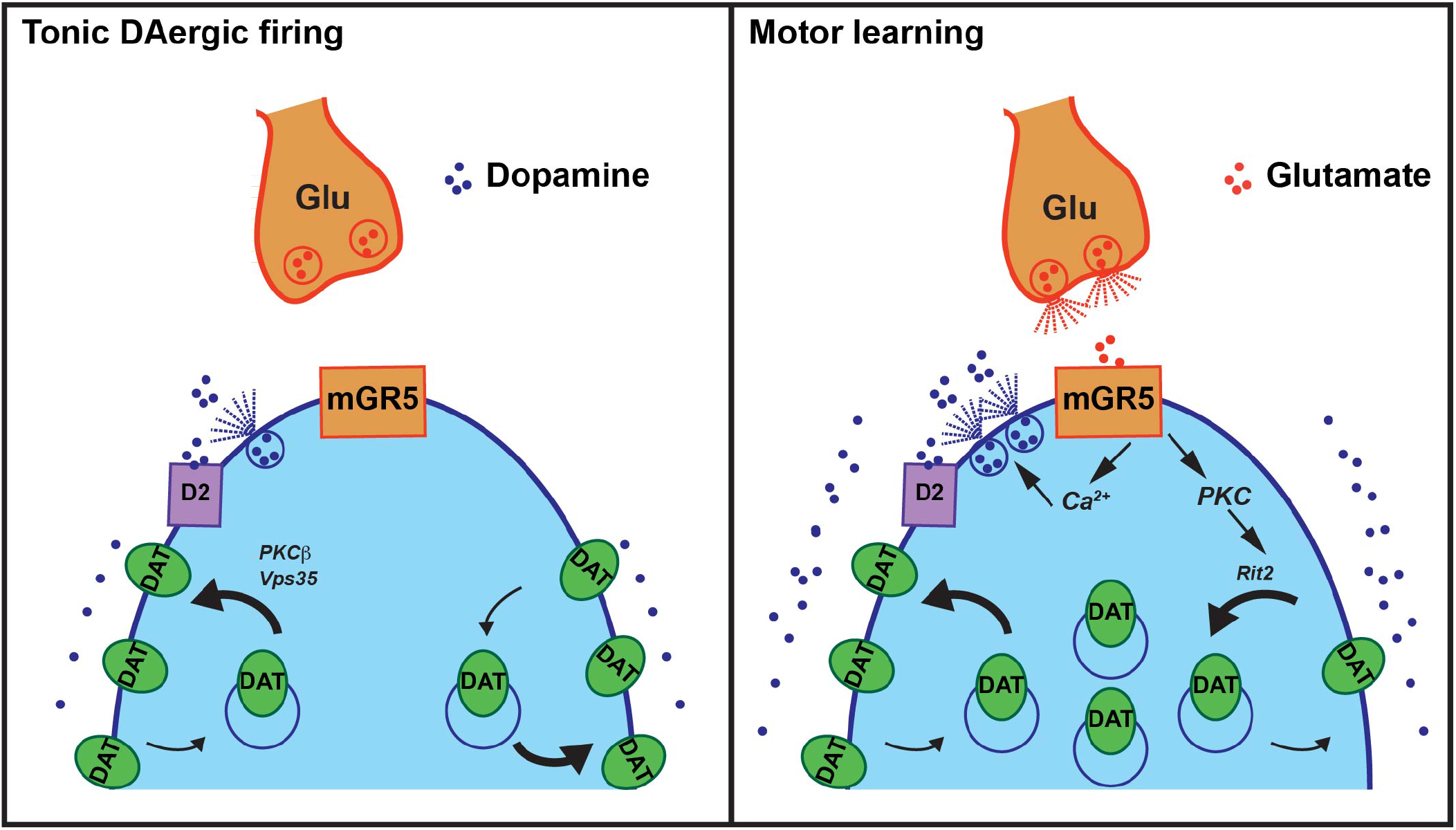
Striatal DAT trafficking model. A presynaptic DAergic terminal and a glutamatergic afferent are depicted. (*Left) Tonic DA neuron firing*. During tonic DA release, DRD2_auto_ activity robustly delivers DAT to the plasma membrane in a PKCb and Vps35-dependent manner. Dense DAT surface expression restricts extracellular DA levels. (*Right) Glutamate release during motor learning.* When DA demand is higher, such as during motor learning, glutamate release from striatal glutamatergic afferents activates presynaptic mGluR5 (mGR5), which elevates intracellular calcium (Ca^2+^) and enhances DA release and DAT membrane insertion. However, mGluR5-mediated PKC activation concurrently drives DAT membrane retrieval in a Rit2-dependent manner. DAT retrieval decreases DAT surface levels, relative to tonic firing, to sustain extracellular DA levels during a period of increased demand.

To date it was unknown whether or how regulated DAT trafficking occurred in DAergic terminals downstream of native presynaptic receptor activation. Moreover, given the well-known interplay within the striatum among DAergic, cholinergic, and glutamatergic release, it was also unclear whether pharmacologically induced DAT trafficking was mediated by cell autonomous mechanisms in DAergic terminals, or by cell non-autonomous mechanisms elsewhere within the striatum. In the current study, we leveraged a variety of chemogenetic and pharmacological approaches, coupled with AAV-mediated conditional gene silencing, to determine how DAT surface expression is acutely modulated by presynaptic receptor activation. Moreover, we asked whether receptor-stimulated DAT trafficking impacts either DA signaling in the DS or motor function. Our results reveal that DAT traffics biphasically in response to Gq receptor activation selectively in DAergic terminals, via either hM3Dq (Figs. 1,2) or the presynaptic mGluR5 (Figs. 3, 4). These results were initially surprising, in light of copious reports by our laboratory^24, 25, 27–29, 36, 57, 58^ and others^26, 59–61^ that direct PKC activation decreases DAT surface expression. Moreover, a previous report in transfected cell lines demonstrated that activating the Gq-coupled neurokinin receptor decreased DAT surface expression in a PKC-dependent manner^31^. A more recent study similarly demonstrated that the Gq coupled muscarinic receptors M1 and M5 stimulate DAT internalization in primary DA neuronal culture and in mouse midbrain^30^. Interestingly, our results indicated that PKC-stimulated DAT internalization does, indeed occur, however, in the context of our current findings, it appears that PKC-stimulated DAT internalization is a retrieval mechanism that is engaged following Gq-coupled receptor activation that tempers DAT surface levels following stimulated DAT membrane insertion.

Our findings demonstrated that the initial Gq-stimulated DAT insertion requires DA release and DRD2 activation. Previous studies in synaptosomes and transfected cell lines reported that DAT function and surface expression rapidly increase in response to treatment with quinpirole^40, 62^, a non-selective DRD2/DRD3 agonist. Moreover, DRD3 activation was sufficient to increase DAT surface expression in non-neuronal cell lines co-transfected with DRD3 and DAT^63^. Both DRD2 and DRD3 are expressed in the striatum^64^, and heretofore it was not known whether quinpirole-stimulated DAT trafficking in terminals was mediated by DRD2, DRD3, or both. Moreover, it is also unknown whether DRD2 activation impacts DAT directly in presynaptic terminals, or via indirect means within the striatum where DRD2 is widely expressed and modulates both glutamate^65–67^ and acetylcholine^68^ release. We found that Gq-stimulated DAT insertion, driven by either hM3Dq (Fig. 2) or mGluR5 (Figs. 4,5), selectively requires presynaptic DRD2_auto_ silencing. Thus, our results suggest that 1) presynaptic DRD2 activation is sufficient and required for Gq-stimulated DAT insertion, and 2) DRD2 activation elsewhere within the striatum does not contribute to DAT membrane insertion in response to Gq-evoked DA release.

Increasing evidence suggests that DA homeostasis and DAT regulation may differ regionally in VS vs. DS^69^. We also observed region-specific differences for DRD2-and Gq-stimulated DAT trafficking. Direct DRD2 activation with sumanirole was sufficient for DAT membrane insertion in both DS and VS (Fig. S1). However, DAT remained elevated at the plasma membrane following DRD2-stimulated membrane insertion in the DS, whereas DAT returned to baseline in the VS, despite lack of any exogenous Gq receptor activation (Fig. S1b). Moreover, following Gq-stimulated DAT insertion, we observed differential kinetics for PKC-dependent retrieval in VS vs. DS, with significantly faster retrieval in VS vs. DS (Fig 1). It is currently unknown whether DAT and/or DRD2 levels or expression ratio differ in VS and DS, and it is possible that their differential expression could contribute to these regional differences. There may also be differential spontaneous glutamate release from glutamatergic terminals in VS vs. DS, which would drive DAT retrieval via mGluR5 in VS preferentially to DS. It is also possible that glutamate co-release from DA terminals may play an autocrine regulatory role in VS, a hypothesis supported by optogenetic studies which demonstrate that glutamate co-release from DA neurons occurs in VS, but not DS^70, 71^. Another possible contributor to striatal DAT trafficking could arise from striatal cholinergic interneurons signaling via the Gq-coupled M5 receptor, which drives DAT internalization in midbrain^30^ and is reportedly expressed in striatal DAergic terminals where it potentiates DA signaling^72^. Future studies will explore how glutamatergic and cholinergic signaling converge onto DA terminals to influence DAT surface levels and DA signaling.

We previously reported that the neuronal, ras-like GTPase, Rit2, is required for PKC-stimulated DAT endocytosis in both cell lines^27, 29^ and DAergic terminals^29^ in response to phorbol ester treatment. We additionally found that endocytosed DAT primarily targets to retromer^+^ endosomes, and that intact retromer complex is required for DAT membrane delivery in neuroblastoma cell lines^28^. However, it was unknown whether mechanisms required for DAT internalization and membrane delivery identified in cell lines, or in response to direct PKC activation, would likewise be required for Gq-stimulated, biphasic DAT trafficking in DAergic terminals. Using AAV-mediated, conditional gene silencing to deplete either Vps35 or Rit2 in DA neurons, we determined that that Vps35^+^ retromer complex is required for stimulated DAT membrane insertion (Fig. 5), and that Rit2 is required for PKC-dependent DAT retrieval, but not PKCβ-mediated DAT insertion (Fig. 6). Consistent with our findings, a recent study found that DAT and Vps35 co-localize in axons of the medial forebrain bundle^73^. Interestingly, apart from their roles in DAT trafficking (and other cellular processes), both Vps35 and Rit2 have been identified as PD risk factors. A Vps35 mutation (D620N) from late onset PD patients was recently reported^74^, and a D620N-Vps35 mouse model exhibited reduced DAT surface expression^75^, consistent with disrupted, retromer-dependent DAT membrane delivery. Rit2 has also been linked to PD in multiple GWAS studies^76–81^, further highlighting its importance in DA signalling and DA neuron function. However, whether dysregulated DAT trafficking contributes to the progressive PD phenotype is currently unknown.

How does biphasic DAT trafficking impact DA signaling and/or DA-dependent behaviors? To begin to address this challenging question, we conditionally silenced mGluR5 in *Pitx3^IRES-tTA^;mGluR5^fl/fl^* midbrain DA neurons. Although mGluR5 is expressed at high levels in the striatum, it was not known whether mGluR5 is expressed presynaptically in DAergic terminals. Conditional mGluR5 silencing completely abolished DHPG-stimulated DAT membrane insertion (Fig. 4c), clearly demonstrating that mGluR5 is expressed in DA terminals, where it directly drives biphasic DAT trafficking. Furthermore, DAergic mGluR5 silencing increased basal DAT surface expression (Fig. 4d) in both VS and DS, suggesting that mGluR5 is specifically pivotal for setting DAT surface tone.

We explored the impact of both DRD2-and mGluR5-stimulated DAT trafficking on DA release and clearance in DS by FSCV. Given that DAT mediates DA clearance and that clearance is dependent on the amount of DAT present at the membrane, we focused on monitoring the decay constant, tau, as a readout for changes in clearance due to DAT trafficking. Electrically evoking DAT release in ACSF, will also drive release of multiple striatal neurotransmitters close to the recording site, including acetylcholine, glutamate, and neuropeptides. Therefore, the tau measured under ACSF conditions likely reflects the summated DAT trafficking induced via multiple signaling pathways including DRD2-mediated insertion and mGluR5 mediated retrieval. DRD2 inhibition significantly lengthened DA clearance times in slices from control mice (Fig. 7d), consistent with lower DAT surface levels in the absence of any DRD2 activation. We hypothesized that since conditional mGluR5 silencing increased basal DAT surface expression, this would translate into more rapid clearance rates in ASCF, as DAT would be inserted via DRD2 activation, but would fail to be rapidly retrieved in the absence of presynaptic mGluR5. To our surprise, DAergic mGluR5 silencing significantly increased the tau to clear DA under basal conditions as compared to control mice (Table I), which suggests a potential for suppressed intrinsic DAT function. Moreover, while DRD2 inhibition increased the clearance tau under basal conditions, it had no further effect on DA clearance rates in the absence of mGluR5 (Fig. 7h and Table I), suggesting that mGluR5 may impacts DRD2 function. Indeed, the DA transient amplitude was also unaffected by DRD2 inhibition following presynaptic mGluR5 silencing, as compared to controls (Fig. 7g), raising the possibility that mGluR5 deletion may impact DA release *in vivo,* either via modulating voltage-gated Ca2+ channels, or via Kv1 channels, which are reported to influence DA release^82, 83^. Alternatively, several reports indicate that DA receptors including DRD2 and DRD3 undergo PKC-stimulated internalization^84, 85^. It is possible that, in addition to regulating DAT surface expression, mGluR5 may regulate DRD2 and/or DRD3 surface expression. Although our data suggest that DRD2 blockade alone is sufficient to block mGluR5 mediated DAT insertion, we cannot rule out that in the absence of mGluR5 there may be an aberrant DRD3 contribution that modulates DAT surface levels during evoked DA release. Indeed, DRD2 and DRD3 have been shown to differentially regulate DAT function^86^. While we did observe increased DAT surface expression in quiescent slices, it is possible that other signaling cascades, activated by electrically evoked neurotransmitter release may integrated to suppress DAT surface expression and/or intrinsic function, even in the absence of DAergic mGluR5.

Although DAergic mGluR5 silencing had little effect on baseline locomotor behavior, it significantly disrupted motor learning and coordination measured on the Rotarod (Fig. 8a,b), but had no overall effect on coordinated movement on the challenge balance beam (Fig. 8c-e). A previous study locally applied mGluR5-specific antagonists and found that striatal mGluR5 activity promotes accelerating rotarod performance^87^, consistent with our findings. Conversely, global mGluR5 knockout ameliorated poor rotarod performance in a mouse Huntington’s disease model^88^, suggesting that mGluR5 activation can have different valence effects on motor function, depending on the cell types in which it is expressed. mGluR5 knockout mice also exhibit deficits in spatial learning memory and fear conditioning acquisition^89^, both of which are DAergic dependent learning behaviors, implicating mGluR5 and potentially DAT trafficking in broader learning behaviors. Further studies that utilize *in vivo* voltametric or postsynaptic DA sensors would be pivotal for understanding DA dynamics during complex behaviors.

We hypothesized that presynaptic mGluR5 silencing would disrupt DAT retrieval, leading to increased surface DAT, faster *in vivo* clearance times, and insufficient extracellular DA to facilitate rotarod performance. Consistent with this hypothesis, conditional mGluR5 silencing significantly increased baseline DAT surface expression in VS and DS (Fig. 4d). Moreover, conditional mGluR5 silencing had no effect on total DAT and TH protein levels in either VS or DS (Fig. S3), nor did it impact activated TH, as determined by quantifying pSer40-TH (Fig. S3). Thus, it is possible that DAT trafficking dysregulation contributed to the motor phenotypes we observed. To test this hypothesis, we administered a novel DAT specific inhibitor, CE-158 to mGluR5 silenced mice and DAT inhibition robustly rescued accelerating rotarod performance (Fig. 8g). This strongly suggests that the motor learning deficit is due to the observed increases in DAT surface expression. However, there are several other factors that could underlie and/or contribute to the lack of motor learning following mGluR5 silencing. DAergic mGluR5 silencing could be inhibiting DA neuron excitability at the somatodendritic level and suppressing increased DA neuron firing rates. Consistent with this possibility, chemogenetic Gq-coupled receptor activation in VTA DA neurons was recently demonstrated to increase their firing rates^39^, and activation of Group I mGluRs in midbrain and basal ganglia non-dopaminergic neurons likewise increased neuronal firing^90^. However, although it was recently reported that Group I mGluR activation depresses iPSCs in VTA DA neurons, it appears to do so via mGluR1, not mGluR5, suggesting that mGluR5 may not contribute to DA neuron excitability in VTA^91^. However, it is entirely possible that DAergic mGluR5 silencing may also have impacted either K+ or Ca2+ channel conductances and thereby impacted DA neuron excitability and/or DA release probability in terminals, and that these perturbations contributed to the motor dysfunction we observed. Future studies examining these parameters should be informative in this regard.

In summary, we found that Gq-coupled, presynaptic receptors drive dynamic, biphasic DAT trafficking that differs in VS and DS, and identified the specific presynaptic mechanisms that are required for regulated DAT trafficking *in situ.* Importantly, these studies are the first to demonstrate a cell-autonomous DAT trafficking mechanism driven by an endogenous Gq-coupled GPCR in intact DA terminals, and suggest that glutamatergic signaling onto DAergic terminals may shape DAergic transmission via mGluR5-stimulated DAT trafficking and that DAT trafficking is integral to DAergic behaviors, such as motor learning.

## Methods

### Materials

Clozapine N-oxide (CNO, 4936), reserpine (2742), L-741,626 (1003), LY 333531 (ruboxistaurin, 4738), sumanirole maleate (2773), GF 109203X (BIM I), and (*RS*)-3,5-dihydroxyphenylglycine (DHPG, 0342) were from Tocris. (S,S)-CE-158 was synthesized as previously described^56^. All other reagents were from either Sigma-Aldrich or Fisher Scientific and were of the highest possible grade.

### Mice

*Pitx3^IRES-tTA^/+* mice (on the C57Bl/6J background) were the generous gift of Dr. Huaibin Cai (National Institute on Aging) and were continuously backcrossed to C57Bl/6J mice (Jackson Laboratories). *TRE-hM3Dq* (#014093), *Drd2^fl/fl^* (#020631), *mGluR5^fl/f^* (#028626) mice, all on the C57Bl/6J background, were obtained from Jackson Laboratories and were backcrossed to either C57Bl/6J or to *Pitx3^IRES-tTA^/+* mice to generate *Pitx3^IRES-tTA^*;*TRE-hM3Dq*, *Pitx3^IRES-tTA^*;*Drd2^fl/fl^,* and *Pitx3^IRES-tTA^*; *mGluR5^fl/fl^*mice, respectively. Mice were maintained in 12hr light/dark cycle (lights on at 0700) at constant temperature and humidity. Food and water were available ad libitum. All studies were conducted in accordance with UMass Medical School IACUC Protocol 202100046 (formerly A-1506, H.E.M.).

### AAVs and stereotaxic surgeries

#### AAVs

pscAAV-TRE3g-eGFP and pscAAV-TRE3g-miR33-shRit2-eGFP AAV9 particles were produced as described^35^. <u>pscAAV-TRE3g-shVps35-eGFP and pscAAV-TRE3q-Cre-eGFP</u>: pscAAV-TRE3g-miR33-shRit2-eGFP was digested with BglII and PstI as backbone. miR33-shVps35 insert was synthesized as a gene block with shRNA antisense sequence ATA ATC CAG AAC ATT ACT AAG (V2LMM_36638, Dharmacon), previously validated^54^. Cre recombinase insert was PCR amplified with addition of 5’ BglII and 3’ PstI sites from pAAV-CB6-PI-Cre (UMass Viral Vector Core). High titer AAV9 particles were produced by the University of Massachusetts Medical School Viral Vector Core as previously described^29, 35^.

#### Survival surgeries

Mice aged 3-4weeks were anesthetized with I.P, 100mg/kg ketamine (Vedco Inc.) and 10mg/kg xylazine (Akorn Inc.). 20% mannitol (NeogenVet) was administered I.P. >15 minutes prior to viral delivery, to increase viral spread^92^. Mice were prepared and placed in the stereotaxic frame (Stoelting Inc.). 1ul of the indicated viruses were administered bilaterally to the VTA (Bregman: anterior/posterior: −3.08mm, medial/lateral: ±0.5mm, dorsal/ventral −4.5mm) at a rate of 0.2uL/min. Syringes were left in place for a minimum of 5 minutes post-infusion prior to removal. Viral incubation was a minimum of 4 weeks. Viral expression was confirmed visually by the presence of GFP in the midbrain, and/or by RT-qPCR.

### RNA extraction and RT-qPCR

RNA was isolated from mouse midbrain punches using RNAqueous®-Micro Kit RNA isolation (Thermo Fisher Scientific). Ventral midbrain samples were bilaterally collected from 300µm acute coronal mouse midbrain slices using a 1.0mm^2^ tissue punch. Slices were visualinzed on an inverted fluorescence microscope during punching, to confirm cell transduction via GFP reporter expression, and to enrich for GFP-positive cells. RNA was extracted and reverse transcribed using RETROscript® reverse transcription kit (Themo Fisher Scientific). Quantitative PCR was performed and analyzed using the Applied Biosystems® 7500 Real-Time PCR System Machine and software or using the Bio-Rad C1000 Touch Thermal Cycler with CFX96 Real-Time system and software, using Taqman® gene expression assays for mouse DRD2 exon 2-3 (Mm00438541_m1), Vps35 (Mm00458167_m1), Rit2 (Mm0172749_mH), mGluR5 exon 7-8 (Mm01317985_m1), and GAPDH (Mm99999915_g1).

### Ex vivo slice biotinylation

Surface proteins in acute coronal slices were covalently appended with biotin as previously described by our laboratory^25, 35, 93, 94^. Coronal slices were prepared from 5-9-week old C57Bl/6J and *Pitx3^IRES-tTA^; TRE-HA-hM3Dq* mice (CNO and DHPG studies), or 4-6 weeks following viral injection (Drd2^fl/fl^, shVps35, shRit2, mGluR5^fl/fl^ studies). All data were obtained from a minimum of 3 independent mice, from multiple striatal slices per mouse. Mice were sacrificed by cervical dislocation and rapid decapitation. Heads were immediately submerged in ice-cold NMDG cutting solution, pH 7.4 (20mM HEPES, 2.5mM KCl, 1.25mM NaH_2_PO_4_, 30mM NaHCO_3_, 25mM glucose, 0.5mM CaCl_2_·4H_2_O. 10mM MgSO_4_·7H_2_O, 92mM N-methyl-D-glucamine, 2mM thiourea, 5M Na^+^-ascorbate, 3mM Na^+^-pyruvate). Brains were removed, glued to VT1200S Vibroslicer (Leica) stage and submerged in ice-cold, oxygenated cutting solution. 300µm slices were prepared and slices were hemisected along the midline prior to recovering in ACSF (125mM NaCl, 2.5mM KCl, 1.24mM NaH_2_PO_4_, 26mM NaHCO_3_, 11mM glucose, 2.4mM CaCl_2_·4H_2_O,1.2mM MgCl_2_·6H_2_O, pH 7.4) for 40 min at 31°C. Hemi-slices were treated with the indicated drugs for the indicated times at 37°C with constant oxygenation. Following drug incubations, slices were moved to ice and surface DAT was labeled by biotinylation with the membrane-impermeant sulfo-NHS-SS-biotin as previously described^25, 29, 35, 36, 94^. Striata were further sub-dissected to isolate VS and DS by cutting hemi-slices in a line from the lateral ventricle to lateral olfactory tract (as shown in Fig.1b). Tissue was lysed in RIPA buffer (10mM Tris, pH 7.4; 150mM NaCl; 1.0mM EDTA; 0.1% SDS, 1% Triton X-100, 1% Na deoxycholate) containing protease inhibitors (1.0mM phenylmethylsulfonyl fluoride and 1.0g/mL each leupeptin, aprotinin, and pepstatin) and Phosphatase inhibitor cocktail V (EMD Millipore) was (when evaluating protein phosphorylation). Tissue was disrupted by triturating sequentially through a 200µL pipette tip, 22-and 26-gauge tech-tips and solubilized by rotating (30 min, 4°C). Insoluble material was removed by centrifugation and BCA protein assay (Thermo Fisher Scientific) was used to determine protein concentrations. Biotinylated proteins were quantitatively isolated with streptavidin agarose beads (Thermo) overnight with rotation (4°C), at a ratio of 20µg lysate to 30µL streptavidin agarose beads, empirically determined to be in the linear range of bead saturation for biotinylated striatal proteins. Beads were washed with RIPA buffer (x3) and recovered proteins were eluted in 2x Laemmli DTT sample buffer by rotation (30 min, room temperature). Eluted proteins and their respective lysate inputs were resolved by SDS-Page and proteins were detected and quantified by immunoblot as described below. Surface DAT populations were calculated by normalizing biotinylated DAT signals to the DAT signals from corresponding inputs in a given hemi-slice.

### Immunoblots

Proteins were resolved by SDS-PAGE and proteins were detected and quantified by immunoblotting with the following antibodies: rat anti-DAT (MAB369, Millipore; 1:2000), rabbit anti-TH (AB152, Millipore, 1:10000), rabbit anti-pSer40 TH (AB5935, Millipore, 1:5000). Secondary antibodies conjugated to horseradish peroxidase were all from Jackson ImmunoResearch, and immunoreactive bands were visualized by chemiluminescence using SuperSignal West Dura (Thermo Scientific). Non-saturating immunoreactive bands were detected using a either VersaDoc 5000MP or ChemiDoc imaging stations (Bio-Rad) and were quantified using Quantity One software (Bio-rad).

### Fast-scan cyclic voltammetry

Striatal slices were prepared as described for *ex vivo* slice biotinylation and recovered at 31°C for a minimum of 1 hour prior to recording in oxygenated ASCF supplemented with 500µM Na-Ascorbate. Glass pipettes containing a 7µm carbon-fiber microelectrode were prepared and preconditioned in ASCF by applying triangular voltage ramps (−0.4 to +1.2 and back to −0.4 V at 400 V/s), delivered at 60Hz for 1 hour. Recordings were performed at 10Hz. Electrodes were calibrated to a 1µM DA standard prior to recording. Electrodes were positioned in DS and DA transients were electrically evoked with a 250µA rectangular pulse every 2 min, using a concentric bipolar electrode placed ∼100µm from the carbon fiber electrode. Data were collected with a 3-electrode headstage, using an EPC10 amplifier (Heka) after low-pass filter at 10 kHz and digitized at 100 kHz, using Patchmaster software (Heka). A stable baseline was achieved after evoking six consecutive DA transients, after which experimental data were collected. Each biological replicate is the average of three evoked DA transients/slice, and a minimum of 3 independent mice were used to gather data from the indicated number of slices in each experiment. Data were analyzed in Igor Pro, using the Wavemetrics FSCV plugin (gift of Veronica Alvarez, NIAAA). Peak amplitudes were measured for each individual DA transient, and tau was calculated as 1/e according to the equation: y = y_0_ + A^((x-x)/tau))^

### Mouse behavior

#### Locomotion

Mouse activity was individually assessed in photobeam activity chambers (San Diego Instruments) as previously described^35^. Mice were placed in clean gromet-free cages and horizontal, vertical, and fine movement were measured in 5-minute bins for 90 minutes total. In CE-158 studies, mice were tested over two consecutive days. Each session was comprised of 45min habituation, I.P. injection, and 90min recording. Mice were injected with vehicle (30% Kolliphor EL in sterile saline) on day 1 and the indicated CE-158 dose on day 2. CE-158 was prepared fresh on each day of experimentation.

#### Accelerating and Fixed-Speed Rotarod

Mice were habituated to the behavior room in home cage for > 30min with ambient lighting and the RotaRod unit (UgoBasile 47600) running at 4 RPM. Mice were weighed prior to testing. Accelerating RotaRod: Mice were placed on the rod revolving at constant 4 RPM, and rod speed was then increased from 4 to 40RPM over 5 minutes. Mouse latency to fall was measured over three consecutive trials and was determined by either triggering the strike plate during a fall, or if the mouse made >1 consecutive passive rotation. Fixed speed: mice were placed on the rod moving at the indicated speeds (20, 25, 30, 35, 40, 45 RPM) and were evaluated for two consecutive 60 second trials. Latency to fall was measured or trial was stopped following >1 passive rotation.

#### Challenge/Balance Beam

One-day prior to assay, mice were trained (5 trials) to traverse a 1.0m, step-wise tapered (widths: 35mm, 25mm, 15mm, 5mm) elevated beam (#80306, Lafayette Neuroscience) at an incline of 15°. Training and assay were performed in a dark room with only one light source placed approximately 1.5 feet over the beam origin. A dark box with home-cage bedding was placed at the top of the incline. Mice were acclimated to testing room for >30min with assay set up. On assay day, a challenge grid (custom 3D-printed, Thingiverse #4869650) was placed over the beam and mice traversed the beam in 3 independent trials. Traversals were video captured and scored for foot faults and traversal time, averaged over the first two completed trials or as paired analysis between first and second trial. Animal IDs were double-blinded to both the experimenter and an independent scorer.

#### Grip Strength

Mouse grip strength was measured using the Bioseb Grip Strength Test (BIO-GS3) equipped with mesh grip grid for mice. Mice were suspended by the tail over the mesh and were allowed to grab the mesh with all 4 paws. The mouse was then pulled backwards on the horizontal plane until it released from the mesh. The force applied, just before release, was recorded for 3 consecutive trials and averaged.

#### Gait Analysis

Gait analysis assay was adapted from Wertman, *et al.*^95^. Briefly, mouse fore-paws and hind-paws were dipped in orange and blue non-toxic tempera paint, respectively. The mice were placed in a 10cm x 36cm runway with 14cm high foamboard walls and a dark box at the opposing end. Fresh, legal-size paper was placed on the bench top under the runway for each trial and mice were placed on the paper at the open end of the runway and allowed to traverse to the closed box at the opposite end. Three trials were performed per mouse and stride length, stride width, and toe spread were measured for both fore and hindlimbs. Number of completed trials was also quantified. Mouse IDs were double-blinded for the both the assay and quantification.

### Statistical Analysis

Data were analyzed using GraphPad Prism software. All data were assessed for normality and nonparametric tests were applied in the event that data distribution was non-Gaussian. Outliers in a given data set were identified using either Grubb’s or Rout’s outlier tests, with α or Q values set at 0.05 or 5%, respectively, and were removed from further analysis. Significant differences between two values were determined using either a one-tailed, two-tailed, or paired Student’s t test, as indicated. Differences amongst more than two conditions were determined using one-way or two-way ANOVA, as appropriate, and significant differences among individual values within the group were determined by post-hoc multiple comparison tests, as described for each experiment. Power analyses were performed using G*Power to determine sufficient sample sizes for behavioral assays given power (1-β) = 0.9, α=0.05, and effect sizes observed in previous and pilot studies. Minimum sample sizes were as follows: accelerating rotarod n=5/group, fixed-speed rotarod n=5/group, total locomotion n=6/group, grip strength n=6/group, balance beam foot slips n=5/group, balance beam traversal time n=6/group, gait analysis n=5/group.

## Acknowledgments

Experiments were designed by P.J.K., H.E.M. and G.E.M., were performed by P.J.K and T.C, and were analyzed by P.J.K and H.M. P.J.K. and H.E.M wrote the manuscript, with editorial suggestions provided by G.E.M. G.L. supplied CE-158 and provided consultation for its use *in vivo* in mice. We thank Drs. Veronica Alvarez, Roland Bock, and Hoon Shin (NIAAA), for generously giving of their time to train and consult with us for the FSCV studies. We also thank Dr. Kensuke Futai for additional technical assistance with all matters electrophysiological, and Dr. Andrew Tapper for insightful discussions. These studies were supported by NIH grants R01DA035224 (H.E.M.) and F31 DA045446 (P.J.K).

## Materials and Correspondence

Any requests for materials described in these studies should be addressed to Dr. Haley E. Melikian, UMASS Medical School.

## Data Availability

All data generated or analyzed during this study are included in this published article and its supplementary information files.

