## Supplemental data and methods for "Presynaptic Gq-coupled receptors drive biphasic dopamine transporter trafficking that modulates dopamine clearance and motor function"

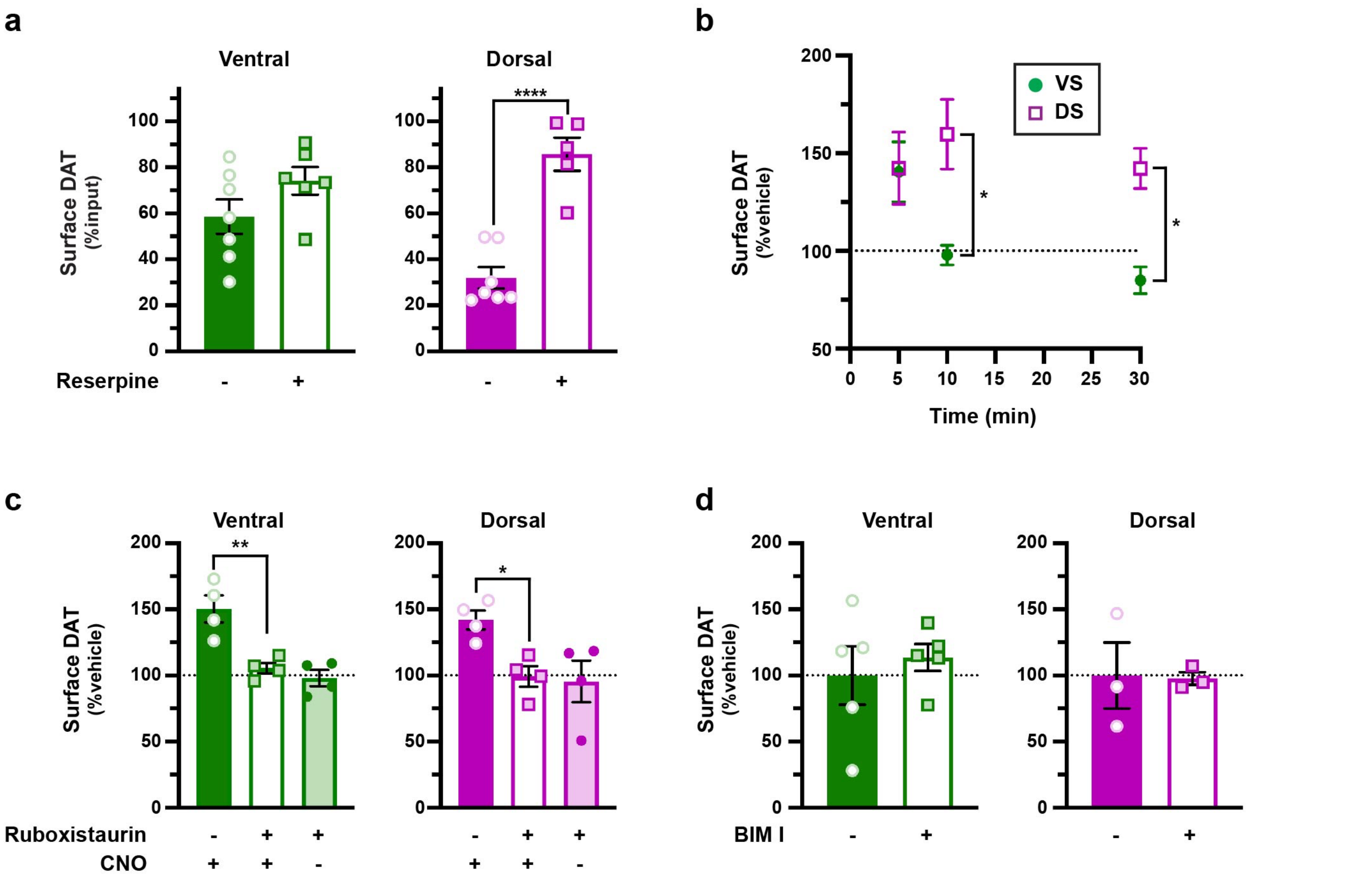


**Figure S1. Mechanisms required for hM3Dq- and DRD2-stimulated DAT membrane delivery and retrieval.** *Ex vivo striatal slice surface biotinylation.* Acute striatal slices were prepared from *Pitx3$^{IRES\text{-}tTA}$;TRE-hM3Dq* mice, were treated with the indicated drugs for the indicated times, and DAT surface levels were measured by slice biotinylation as described in *Methods.* Dorsal and ventral striata were subdissected prior to solubilizing, as described in *Methods.* Mean DAT surface levels are presented ±S.E.M. **(a)** *Effect of reserpine treatment on basal DAT surface expression.* Mice were injected (I.P.) ±5.0 mg/kg reserpine 16 hrs prior to preparing slices, and 1.0μM reserpine or vehicle were included in the bath throughout the experiment. *Ventral:* p=0.14, two-tailed, unpaired Student's t test, n=7 (saline) and 6 (reserpine). *Dorsal:* ****p<0.0001, two-tailed, unpaired Student's t test, n=7 (saline) and 5 (reserpine). **(b)** *DRD2 activation increases DAT surface expression:* Slices were treated with 170nM sumanirole for the indicated times. Two-way ANOVA: Interaction: $F_{(2, 23)}$ =2.92, p = 0.07; Time: $F_{(2, 23)}$ =2.08, p=0.15, Region: $F_{(1, 23)}$ =12.55, **p=0.002. DAT surface expression remained significantly more elevated in dorsal striatum than ventral striatum following 10 min(*p=0.02) and 30 min (p=0.02) treatment times, Bonferroni's multiple comparisons test, n=4-5 (ventral) and 5 (dorsal). **(c)** *hM3Dq-stimulated DAT insertion requires PKC$\beta$ activity.* Slices were pretreated ±ruboxistaurin (50nM, 30 min, 37°C) and DAT insertion was stimulated ±CNO (500nM, 5 min). *Ventral:* One-way ANOVA: $F_{(2,9)}$ =15.10, **p=0.001; Ruboxistaurin significantly blocked CNO-stimulated DAT insertion, **p=0.004, Bonferonni's multiple comparison test, n=4. *Dorsal:* One-way ANOVA: $F_{(2,9)}$ =5.54, *p=0.03; Ruboxistaurin significantly blocked CNO-stimulated DAT insertion, *p=0.04, Bonferonni's multiple comparison test, n=4. **(d)** *Effect of BIM I on DAT surface expression.* BIM I treatment alone had no significant effect on DAT surface levels in either ventral (p=0.55) or dorsal (p=0.93) striata (two-tailed, unpaired Student's t test, n=5 (ventral) and n=3 (dorsal)).

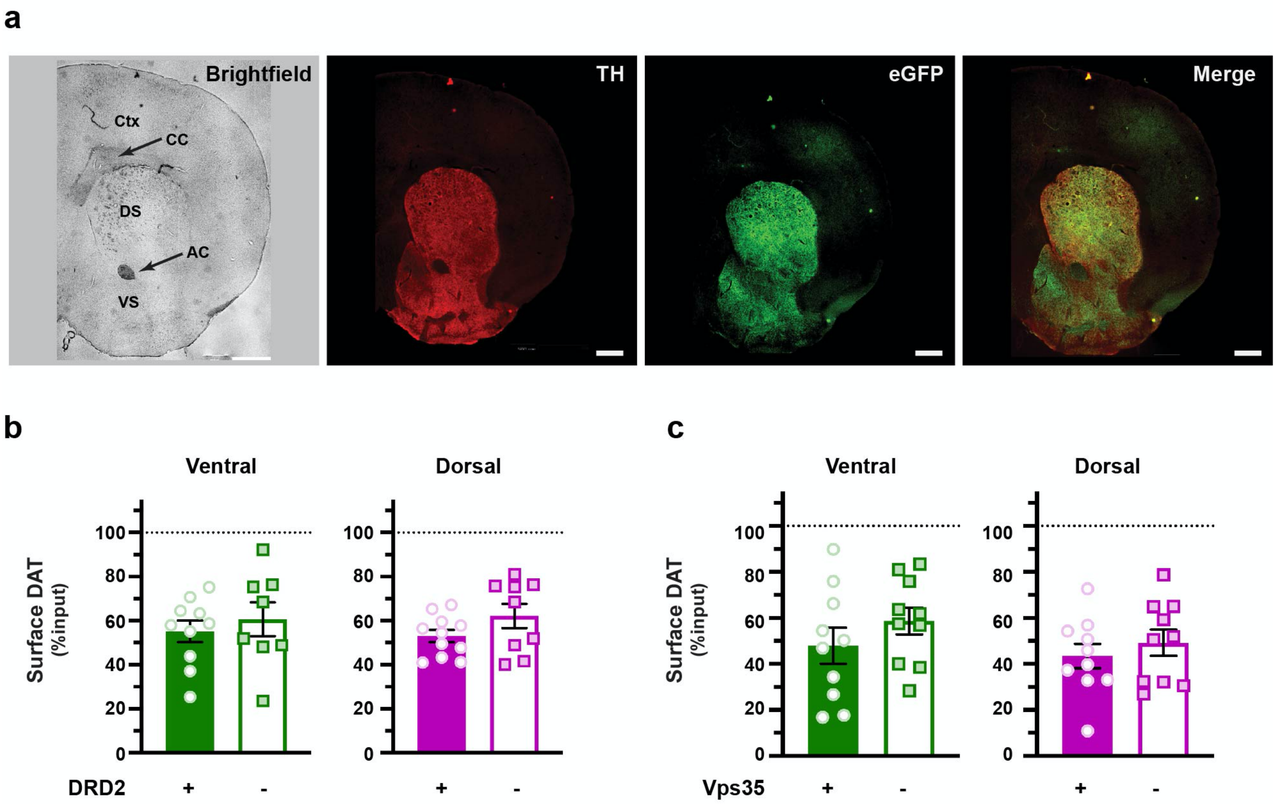


**Figure S2. Effect of conditional DRD2 and Vps35 silencing on basal DAT surface expression.**
**(a)** *AAV9 viral spread validation*. *Pitx3*$^{IRES\text{-}tTA}$ mouse VTA were bilaterally injected with AAV9-TRE-shVPS35-eGFP. Brains were harvested and sectioned (25µm) 4 weeks post-injection, and were stained for TH (red) and eGFP (green) as described in *Methods*. Images were captured at bregma 1.54 in brightfield, red, and green channels as indicated. A representative hemisphere is shown. Note robust eGFP expression in both dorsal and ventral striatum that co-localizes with TH. Ctx (cortex), DS (dorsal striatum), VS (ventral striatum), AC (anterior commissure), CC (corpus collosum). Scale bar = 500µm. **(b, c)** *Ex vivo striatal slice surface biotinylation*. Acute striatal slices were prepared from the indicated mice, and DAT surface levels were measured by slice biotinylation as described in *Methods*. Mean DAT surface levels in ventral (left) and dorsal (right) striata are presented as %input ±S.E.M. (b) DRD2auto silencing: *Pitx3*$^{IRES\text{-}tTA}$;*DRD2*$^{fl/fl}$ mouse VTA were bilaterally injected with AAV9-TRE-Cre. DRD2auto silencing had no significant effect on basal DAT surface levels in either ventral (p=0.54) or dorsal (p=0.13) striata, as compared to non-injected, *Pitx3*$^{IRES\text{-}tTA}$ controls, two-tailed, unpaired Student's t test, n=8-10 (ventral) and n=9-11 (dorsal). (c) Vps35 silencing. *Pitx3*$^{IRES\text{-}tTA}$ mouse VTA were bilaterally injected with either AAV9-TRE-eGFP or AAV9-TRE-shVps35. Conditional Vps35 silencing had no significant effect on basal DAT surface levels in either ventral (p=0.28) or dorsal (p=0.46) striata, as compared to control-injected mice (two-tailed, unpaired Student's t test, n=10 (ventral) and n=10 (dorsal)).

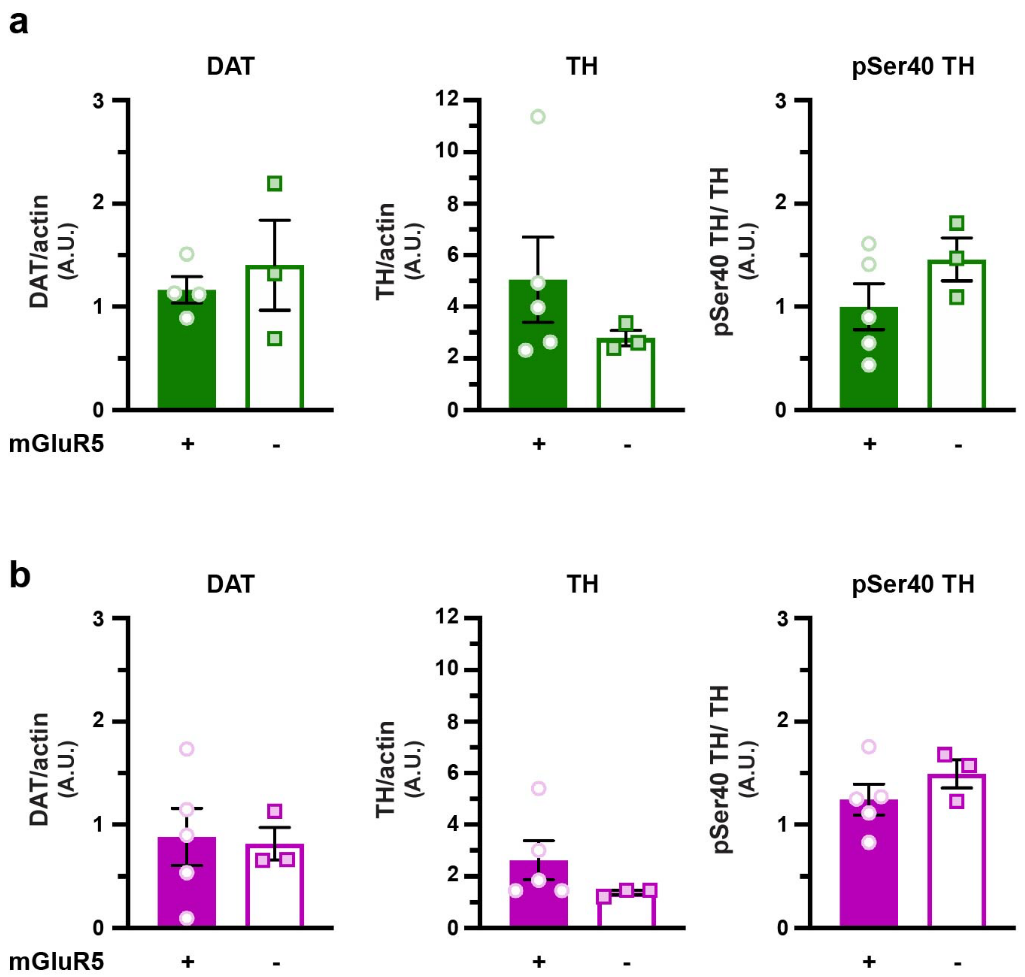


**Figure S3. Effect of conditional mGluR5 silencing on DAT and TH expression and TH activity.** *Quantitative immunoblotting from ventral and dorsal striata. Pitx3$^{IRES\text{-}tTA}$;mGluR5$^{fl/fl}$* mouse VTA were bilaterally injected with either AAV9-TRE-eGFP or AAV9-TRE-Cre and DAT, TH, and pSer40-TH levels were measured in lysates from ventral and dorsal striata. Mean protein levels are presented ±S.E.M., normalized to either actin loading controls (DAT and TH) or pan-TH (pSer40-TH) **(a)** *Ventral striatum:* Conditional mGluR5 silencing had no significant effect on either total DAT (p=0.57, n=4 [eGFP) and 3 [Cre]), total TH (p=0.38, n=5 [eGFP) and 3 [Cre]), or pSer40-TH (p=0.22, n=5 [eGFP) and 3 [Cre]), two-tailed, unpaired, Student t test. **(b)** *Dorsal striatum:* Conditional mGluR5 silencing had no significant effect on either total DAT (p=0.87), total TH (p=0.35), or pSer40-TH (p=0.31), two-tailed, unpaired, Student t test, n= n=5 [eGFP) and 3 [Cre].

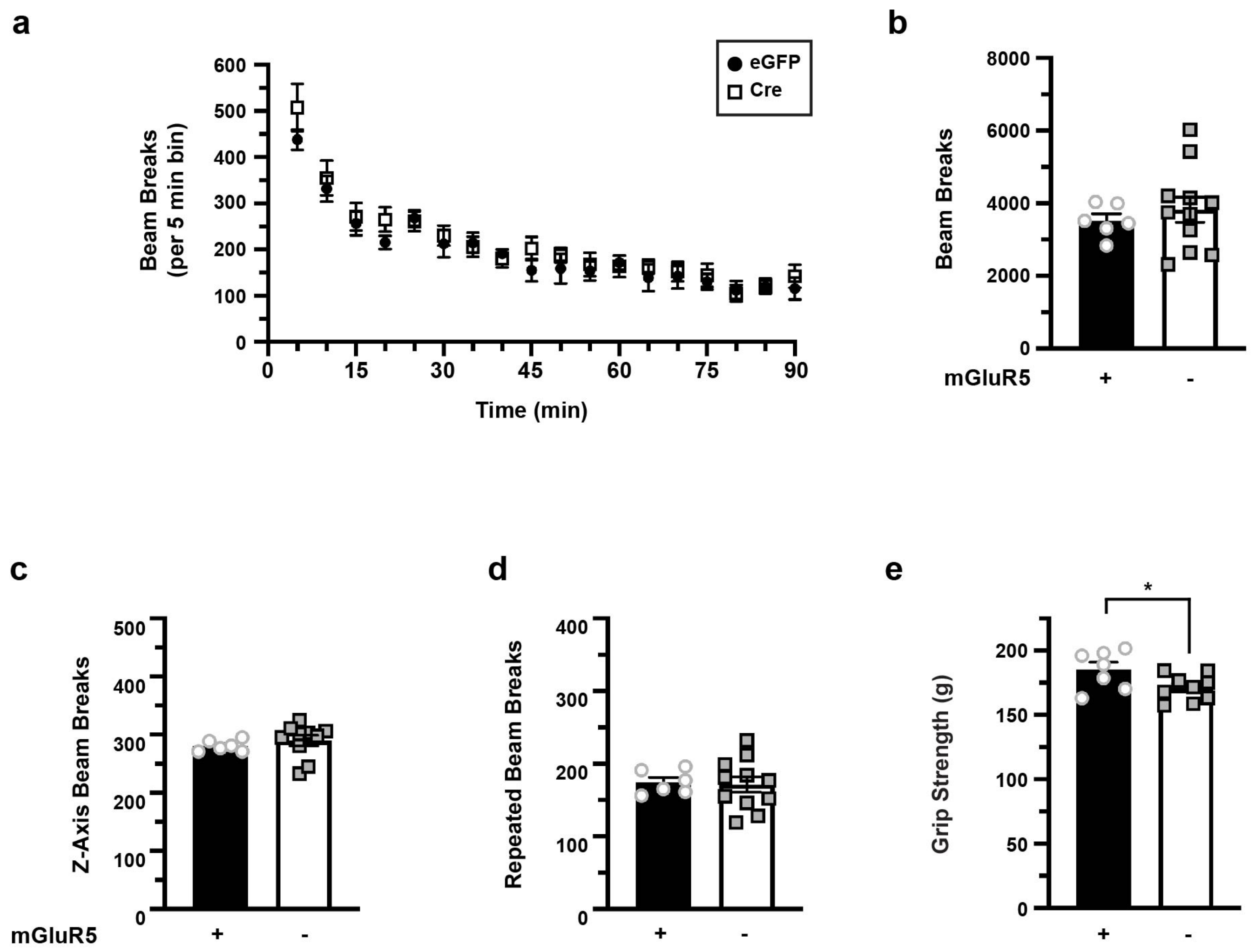


**Figure S4. Effect of conditional mGluR5 silencing on mouse baseline locomotion and grip strength.** *Mouse locomotor studies. Pitx3$^{IRES\text{-}tTA}$;mGluR5$^{fl/fl}$* mouse VTA were bilaterally injected with either AAV9-TRE-eGFP or AAV9-TRE-Cre and baseline locomotor activity was monitored in photobeam activity chambers as described in *Methods.* **(a)** *Total horizontal locomotion over time*. **(b-d)** *Averaged data.* Average total movement measured throughout the recording session, ±S.E.M. Conditional mGluR5 silencing had no significant effect on horizontal locomotion (p=0.55, **b**), vertical movement (p=0.95, **c**), or fine movement (p=0.85, **d**), two-tailed, unpaired, Student t test, n=6 (eGFP) and n=11 (Cre). **(e)** *Grip Strength:* Mouse grip strength was assessed as described in *Methods.* Average data are presented as force required (g) to drive mouse release ±S.E.M. Conditional mGluR5 silencing significantly decreased mouse grip strength (p=0.04), two-tailed, unpaired, Student t test, n=6 (eGFP) and n=11 (Cre).

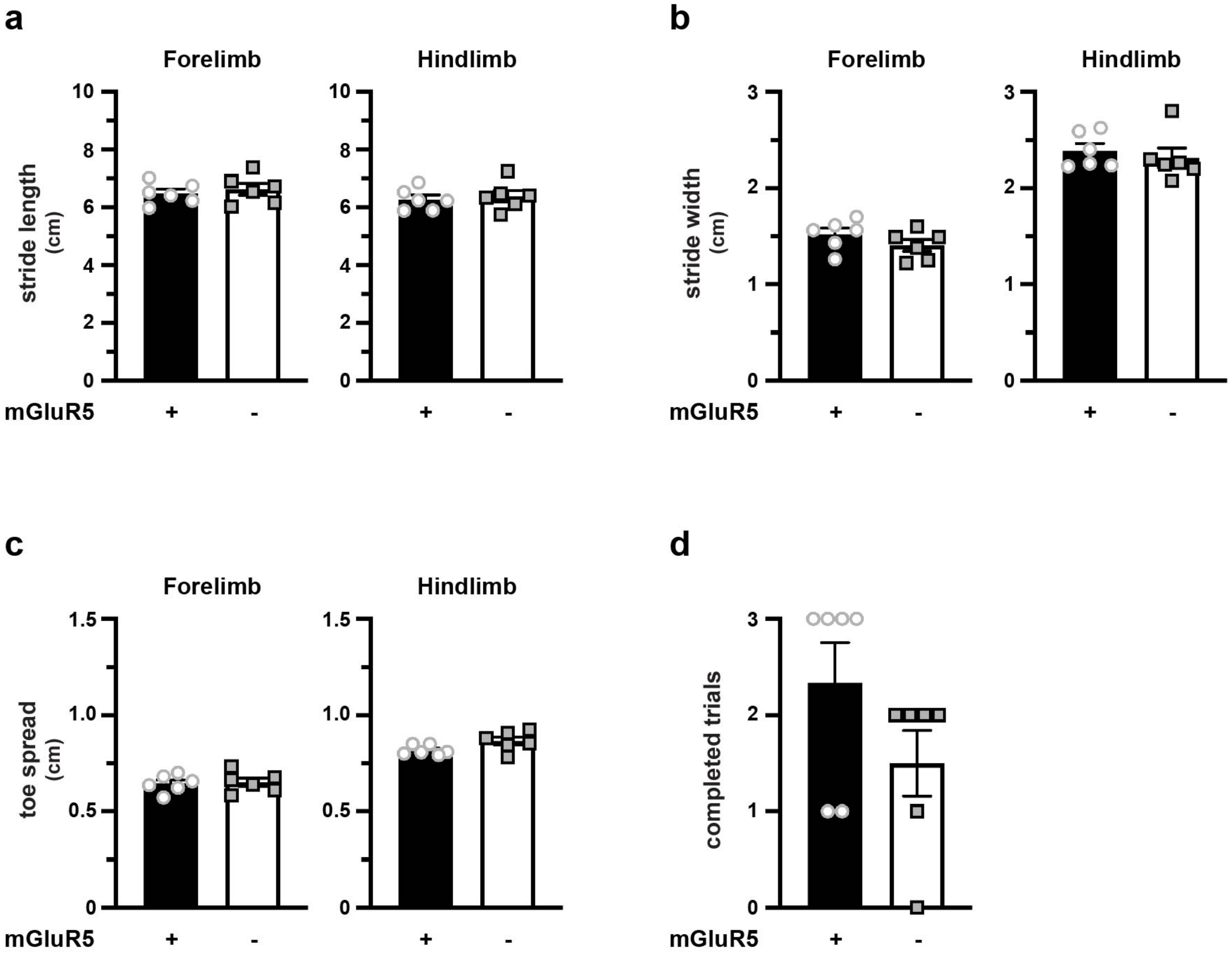


**Figure S5. Effect of conditional mGluR5 silencing on mouse gait.**
*Pitx3*$^{IRES\text{-}tTA}$*;mGluR5*$^{fl/fl}$ mouse VTA were bilaterally injected with either AAV9-TRE-eGFP or AAV9-TRE-Cre and mouse gait was assessed as described in *Methods.* Conditional mGluR5 silencing had no significant effect on either **(a)** stride length in the forelimb (p=0.60) or hindlimb (p=0.66), **(b)** stride width in the forelimb (p=0.21) or hindlimb (p=0.56), **(c)** toe spread in the forelimb (p=0.83) or hindlimb (p=0.06), or **(d)** total completed trials (p=0.16) (two-tailed, unpaired, Student t test, n=6 (eGFP) and n=11 (Cre).

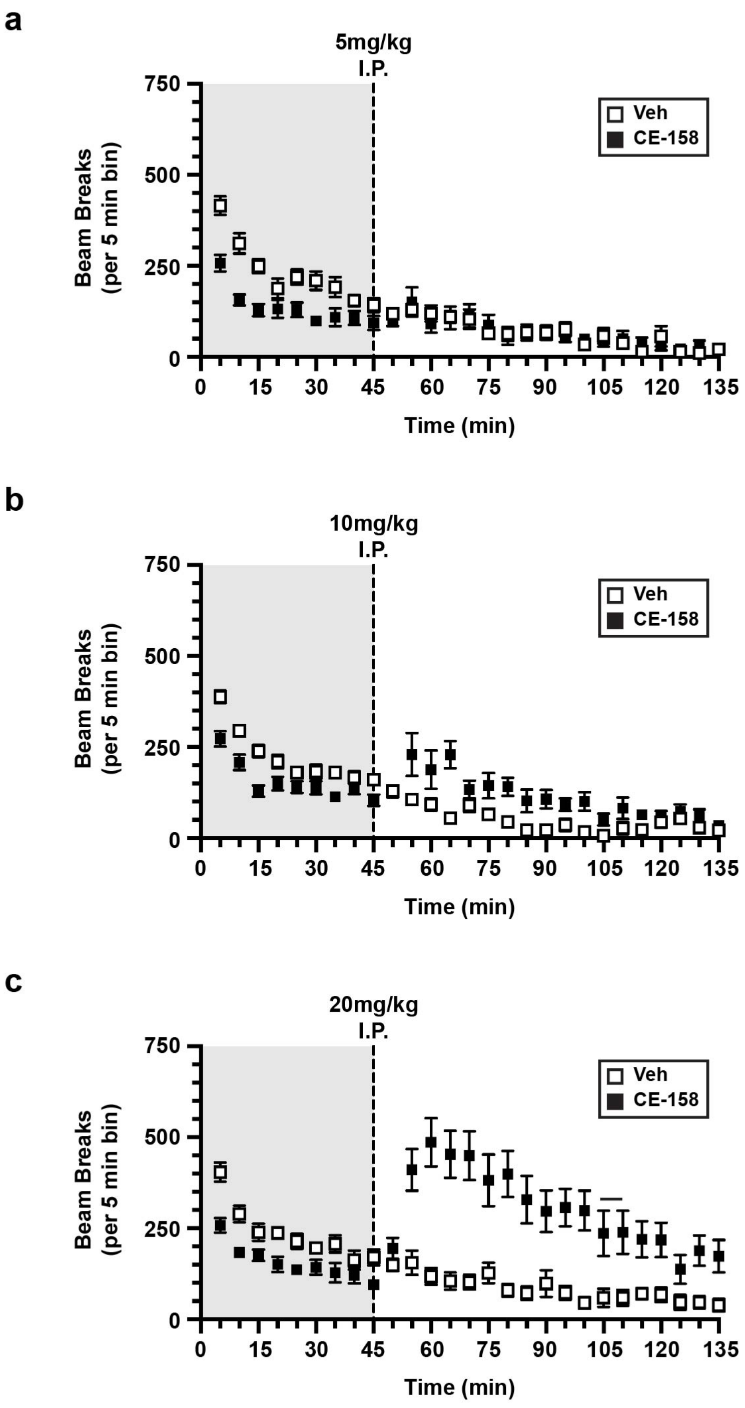


**Figure S6. Effect of DAT inhibitor CE-158 on wildtype mouse horizontal locomotion.**
*Mouse locomotor studies. WT* mouse horizontal locomotor activity was monitored in photobeam activity chambers as described in *Methods.* Mice were habituated to the test cage for 45 min before I.P. injection ±CE-158. Horizontal locomotion was measured over the subsequent 90min. Vehicle and CE-158 data were collected from the same animals over 2 consecutive days of testing. **(a)** *5mg/kg* no significant effect of drug. Two-way Repeated Measures ANOVA: Interaction: $F_{(26, 468)}$ =5.322, ****p < 0.0001; Time: $F_{(26, 468)}$ =39.66, ****p < 00001; Drug: $F_{(1, 18)}$ =3.208, p=0.0901; Subject: $F_{(18, 468)}$ = 15.59, ****p < 0.001; n=10 **(b)** *10mg/kg* no significant effect of drug. Two-way Repeated Measures ANOVA: Interaction: $F_{(26, 468)}$ =9.648, ****p < 0.0001; Time: $F_{(26, 468)}$ =36.02, ****p < 00001; Drug: $F_{(1, 18)}$ =1.057, p=0.3176; Subject: $F_{(18, 468)}$ = 16.74, ****p < 0.001; n=10. **(c)** *20mg/kg* significantly increases horizontal locomotion. Two-way Repeated Measures ANOVA: Interaction: $F_{(26, 468)}$ =15.98, ****p < 0.0001; Time: $F_{(26, 468)}$ =9.570, ****p < 00001; Drug: $F_{(1, 18)}$ =3.208, **p=0.0026.18; Subject: $F_{(18, 468)}$ = 18.74, ****p < 0.001; n=10.
